# The punctate localisation of the yeast sterol transporter Ysp2p is determined by three dimerisation interfaces in its C-terminus

**DOI:** 10.1101/2023.08.08.552482

**Authors:** Ganiyu O. Alli-Balogun, Lazar Ivanović, Wanda Kukulski, Tim P. Levine

**Affiliations:** UCL Institute of Ophthalmology, 11-43 Bath Street, London EC1V 9EL, UK; Institute of Biochemistry and Molecular Medicine, University of Bern, Bühlstrasse 28, 3012 Bern, Switzerland; Graduate School for Cellular and Biomedical Sciences, University of Bern, Mittelstrasse 43, 3012 Bern, Switzerland

**Keywords:** Ergosterol, cholesterol, Alphafold2, transport, membrane, SCF beta-TrCP, self-organisation, oligomerisation

## Abstract

Sterol lipids traffic between intracellular compartments by vesicular and non-vesicular routes. Sterol traffic from the plasma membrane to the endoplasmic reticulum (ER), so-called retrograde traffic, particularly depends on a non-vesicular mechanism, being transported by the ubiquitous family of <u>L</u>ipid transfer proteins <u>A</u>nchored at <u>M</u>embrane contact sites (LAMs, also called GRAMD1/Asters in humans, VASt in plants). LAMs are similar to many lipid transfer proteins in that they localise to membrane contact sites and carry lipids between two organelles. In yeast, the major LAM active at ER-plasma membrane contact sites is Ysp2p, which has a uniquely punctate distribution in the cortical ER. Here, we have comprehensively dissected how Ysp2p achieves its distinctive punctate localisation. We show that the PH^GRAM^ domain of Ysp2p has membrane binding properties similar to its human counterpart GRAMD1B, but that this is not important for punctate localisation of Ysp2p. Instead, all regions necessary for the punctate localisation of Ysp2p at membrane contacts are present in ∼200 residues at the C-terminus of Ysp2p, with a critical region being a small β-sheet that we predict homodimerises. We also study the role of punctate localisation of Ysp2 in its function in retrograde sterol traffic, and show that function does not require the punctate localisation, but instead requires a polybasic region adjacent to the sterol transfer domain. Finally, to investigate the interaction of the polybasic region with the plasma membrane, we examine contacts populated by the Ysp2 C-terminus by electron tomography, and find that they consist of generic cortical ER.

## Introduction

Maintaining the distribution of membrane sterols is a vital cellular function, as shown by the major human diseases that arise from dysregulated sterol distribution, from atherosclerosis to cancer and neurodegeneration (Goldstein and Brown, 2015; Huang et al., 2020; Vance, 2012). Eukaryotic cells synthesise sterol in the endoplasmic reticulum (ER), which then can traffic to other organelles such as the plasma membrane (PM) either in vesicles or by the action of lipid transfer proteins (LTPs). Sterol can then traffic by both vesicular and non-vesicular routes between organelles, with traffic back from the PM to the ER (called retrograde traffic) being a major route for sensing and storing excess sterol (Maxfield and Wustner, 2002). LTPs involved in sterol transport include members of the Steroidogenic Acute Regulatory Transfer (StART) family. For example, in human cells StART-domain 1 (StARD1) (aka StAR) delivers cholesterol to mitochondria (Clark et al., 1994), and StARD4 exchanges cholesterol between PM and endosomes (Iaea et al., 2017).

LTPs are particularly important for retrograde sterol traffic (Kennelly and Tontonoz, 2023). A major class of LTPs active in this role are the <u>L</u>TPs <u>A</u>nchored at <u>M</u>embrane contact sites (LAMs, also known as GRAMD1, Aster or VASt proteins) (Gatta et al., 2015; Khafif et al., 2014; Marek et al., 2020; Naito et al., 2019; Qiu et al., 2023; Sandhu et al., 2018; Trinh et al., 2020; Trinh et al., 2022) are integral ER membrane proteins containing lipid transfer domains specific for sterols (Gatta et al., 2015; Marek et al., 2020; Murley et al., 2015; Sandhu et al., 2018; Zhu et al., 2021) that are distantly related to StART, hence in the wider StARkin domain superfamily (Wong and Levine, 2016). LAM proteins have a characteristic domain architecture (Figure 1A). Starting at the N-terminus, LAMs have intrinsically disordered regions, similar to other LTPs (Jamecna and Antonny, 2021), that also contains regulatory motifs (Roelants et al., 2018). Next is a pleckstrin homology (PH)-like domain of the GRAM type (PH^GRAM^) (Doerks et al., 2000), which can target membranes via either a protein partner (Murley et al., 2015) or lipid partners; in GRAMD1B the PH^GRAM^ binds cholesterol in combination with phosphatidylserine (PS) (Sandhu et al., 2018). The next domain is the sterol-specific StARkin. Finally, there is the region described here as the C-terminus, which in almost all LAMs contains a transmembrane helix (TMH) that embeds the protein in the ER and keeps the StARkin domain in close proximity to the ER at a membrane contact site (Wong and Levine, 2016).

**Figure 1:**
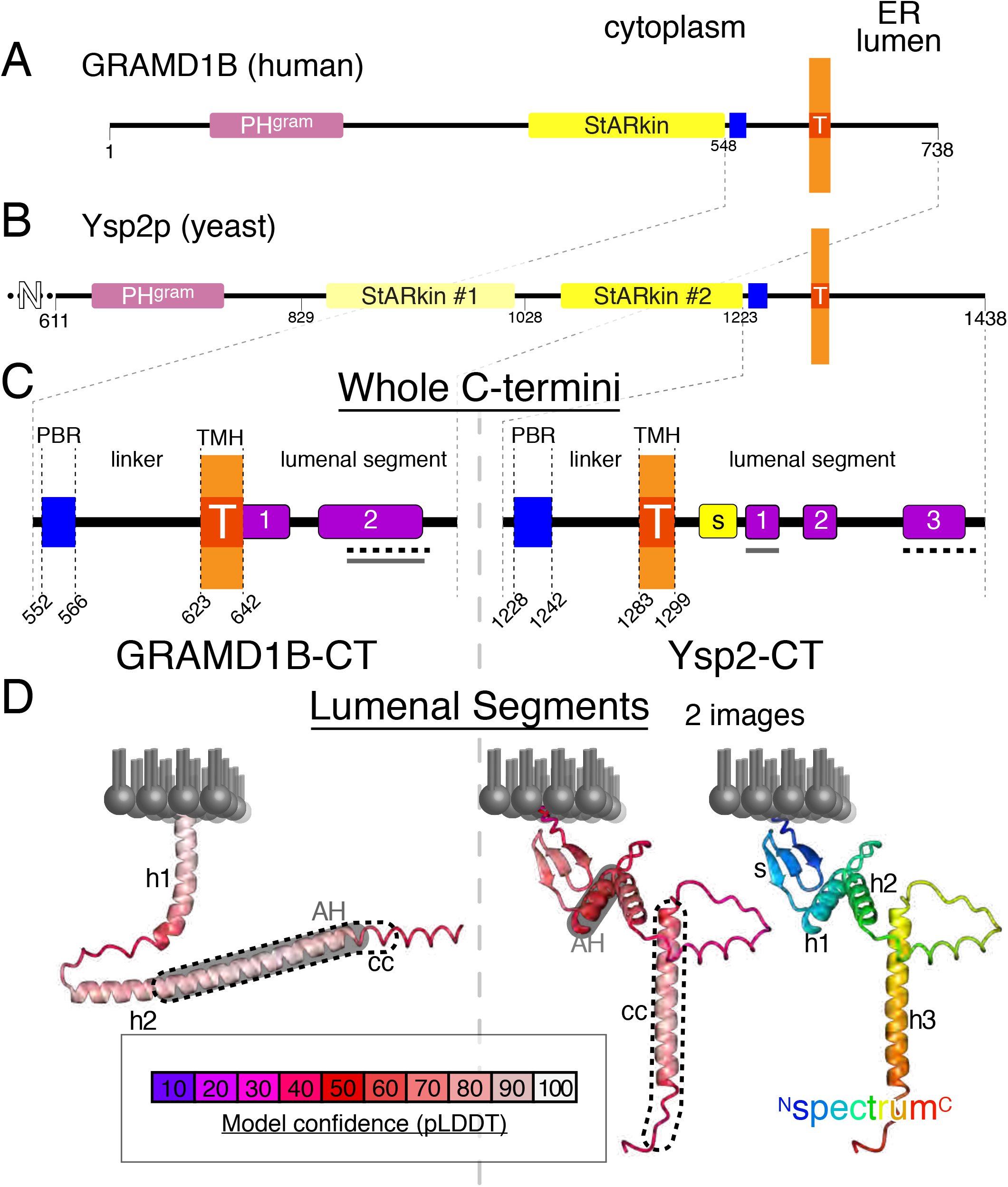
Comparing LAM protein domains, C-termini and lumenal regions from human to yeast A and B. Globular domains in GRAMD1B (human, A) and Ysp2p (*S. cerevisiae*, B). The 2 types of domains present are: PH-like domain of the GRAM family (PH^GRAM^: pink) and sterol-specific StARkin domains (also called StART-like, Aster or VASt domains: yellow; in two shades for the two domains in Ysp2). Also shown are transmembrane helices (TMH: red, marked “T” embedded in the ER membrane, which his orange), and polybasic regions (PBR: blue), which are present at the N-terminal end of the linker between StARkin domain and TMH. The first 600 residues of Ysp2p have no predicted secondary structural elements and have been excluded. Note that Ysp2p has alternate names Ltc4p and Lam2p. **C.** Details of the C-terminal portions of GRAMD1B (192 aa) and Ysp2p (216 aa) that follow the StARkin domains. The C-termini are divided into 4 regions: PBR, linker, TMH and lumenal region. Numbers below indicate the initial residue for each region (except extreme C-terminus). Predicted secondary structural elements in the lumenal regions are shown, β-sheet: yellow, α-helix: purple. Amphipathic helices and predicted coiled coils are indicated by grey and dashed black lines respectively. See Supplementary Figure 3A-E for details of helices and coils. **D.** Structures of lumenal regions of GRAMD1B (left) and Ysp2 (right) predicted by Alphafold (Jumper et al., 2021), shown in relation to the ER membrane, and coloured by model confidence in terms of probability local distance difference test (pLDDT, see key). For Ysp2CT, a second image (far right) is coloured in a blueèred spectrum from NèC-terminus to show the order of elements. Regions with a grey background are amphipathic helices (AH) and regions outlined in dashed black lines are predicted coiled-coils (cc). In GRAMD1B these overlap extensively. Two independent methods predicted the same structural elements: PSIPRED (part C) and AlphaFold (part D) (Buchan et al., 2013; Jumper et al., 2021).

Budding yeast has six LAMs, consisting of three pairs formed by a recent genome duplication (Wolfe and Shields, 1997). Lam5p/Lam6p are most similar to human GRAMD1/Aster proteins, and they localise to ER contacts with mitochondria, vacuoles (equivalent of lysosome) and Golgi compartments (Elbaz-Alon et al., 2014; Murley et al., 2015; Weill et al., 2018). The other two yeast LAM pairs (Ysp2p/Lam4p and Lam1p/Sip3p) are localised to ER-PM contacts. Ysp2p is the main transfer protein for retrograde transport, with Lam4p having only a minor role possibly because of lower expression (Gatta et al., 2015; Murley et al., 2017). By contrast, Sip3p/Lam1p are evolutionarily distant to other LAMs, and have an indirect role in sterol traffic through supporting Ysp2p function (Gatta et al., 2015; Murley et al., 2017).

Ysp2p/Lam4p differ from other LAMs by containing both a second StARkin domain further from the TMH that is not required for function (Gatta et al., 2015) and a highly elongated N-terminus (Figure 1B) (Jamecna and Antonny, 2021). LAMs at the ER-PM interface mediate retrograde sterol traffic in various fungi and animal cells (Chauhan and Fairn, 2021; Gatta et al., 2015; Marek et al., 2020; Sandhu et al., 2018; Trinh et al., 2022; Zhu et al., 2021). In yeast, Ysp2p is found together with Lam4p, Lam1p and Sip3p in punctate ER-PM contacts (Gatta et al., 2015; Murley et al., 2017), which form independently of other ER-PM contact site proteins (Gatta et al., 2015; Manford et al., 2012). In both yeast and human LAM-positive ER-PM contacts do not contain tricalbins/E-Syts cells, members of another LTP family that targets punctate ER-PM contacts (Besprozvannaya et al., 2018; Gatta et al., 2015). Thus, LAM-positive contacts are distinct from other ER-PM contacts. The way human GRAMD1B forms puncta has been studied previously (Naito et al., 2019), but no parallel yeast study has been reported.

Here we have studied how Ysp2p acquires its unique punctate distribution at ER-PM contacts (Naito et al., 2019; Sandhu et al., 2018) by dissecting different regions and comparing to human GRAMD1B. We then applied our findings to determine the contribution of punctate targeting to the function of full-length Ysp2p, finding that it there is no positive correlation, suggesting that punctate targeting may serve another purpose.

## Results

### 1. Punctate targeting by the C-terminus of Ysp2p is independent of all full-length LAMs

We found previously that Ysp2p extensively colocalizes with its paralog Lam4p and the paralogous LAM pair Lam1p/Sip3p (Gatta et al., 2015). Subsequently, these four LAMs were shown to form complexes (Murley et al., 2017). To test if punctate targeting by the Ysp2 C-terminus (Ysp2CT) arises from interactions with LAM partners, we expressed Ysp2CT tagged with GFP tagged at the N-terminus. This construct contains the 216 residues of Ysp2p that follow its second StARkin domain, consisting of: 6 disordered residues, the PBR (15 residues with 6 lysines and 4 arginines), a disordered linker (40 residues), the TMH (16 residues) and finally the lumenal region (140 residues) (Figure 1C). In wild-type strains, Ysp2CT targeted puncta in the cell periphery, with no internal puncta (Figure 2A), which is distinct from the planar localisation characteristic of many ER proteins. We next expressed Ysp2CT in a strain missing all four of the ER-PM LAMs, *lam1*Δ *ysp2*Δ *sip3*Δ *lam4*Δ (Sokolov et al., 2020). Localisation was still punctate, with similar range of brightness of puncta (Figure 2B). The only difference from wild-type cells was that a minority of puncta were internal, possibly on the nuclear envelope. This shows that formation of puncta by Ysp2CT does not require other LAMs, suggesting that punctate localisation arises from self-organisation, which agrees with a previous study of human GRAMD1B-CT (Naito et al., 2019).

**Figure 2:**
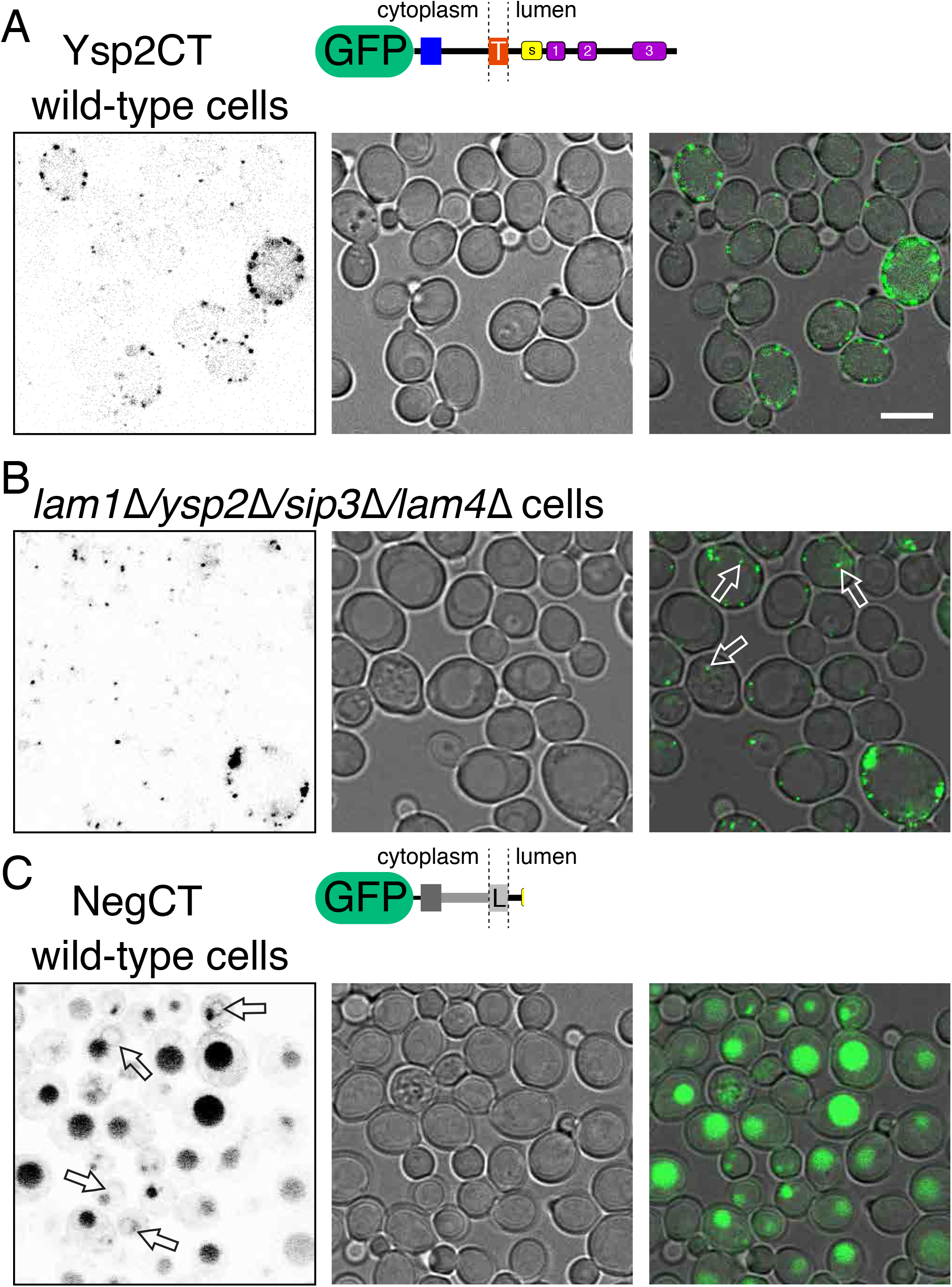
Punctate localisation of Ysp2CT does not require either other LAMs or structural elements previously identified in GRAMD1B. **A.** Intracellular distribution of Ysp2CT (residues 1223-1438), tagged with monomeric Umikinoko Green (here called GFP): LEFT: fluorescence inverted; CENTRE: bright field; RIGHT: merge with fluorescence in green and bright field dimmed by 50%. Constructs (as described in Supplementary Table 1) are based on pRS416, which has an average of 2-5 copies per cell, varying more widely within a single colony (Karim et al., 2013). The merge shows that all fluorescence is peripheral. **B.** GFP-tagged Ysp2CT (as in A) expressed in a *lam1*Δ *ysp2*Δ *sip3*Δ *lam4*Δ strain (Sokolov et al., 2020). Arrows indicate puncta at locations away from the cell cortex. **C.** A negative control construct, made of three generic regions that create the same topology as Ysp2CT, followed by a truncated lumenal region was expressed in wild-type cells. The 89 residues of NegCT after GFP contain only 15 that are from Ysp2. Arrows indicate some of the faint nuclear envelopes. Diffuse fluorescence is inside degradative vacuoles (yeast equivalent of lysosomes). Scale bars = 5 µm.

We next estimated how many molecules are in each punctum by comparing their brightness to puncta of the kinetochore protein Cse4p tagged genomically with the same fluorescent protein. Cse4p forms clusters that in anaphase reach 80 copies per cell (5 per kinetochore), but in asynchronous cultures clusters have 40-50% that number (32-40 molecules) (Coffman et al., 2011; Lawrimore et al., 2011). The fluorescence intensity distribution of the entire population of Cse4-GFP puncta approximately corresponded to the least bright fraction of Ysp2CT puncta, although Ysp2CT showed a range of greater brightness up to 5x this level (Supplementary Figure 1). Thus, we find that Ysp2CT forms puncta with 30-40 copies at minimum.

### 2. The PH^GRAM^ domain of Ysp2 binds PM sterol and is associated with β-TrCP degrons

While constitutive targeting of Ysp2p to ER-PM contacts requires just its C-terminus, human GRAMD1B is located mainly at other contacts deeper inside the cells, for example ER-endosomes (Hoglinger et al., 2019). GRAMD1B only translocates to ER-PM contacts when there is an increase in PM free sterol, which recruits the protein through binding the PH^GRAM^ domain in combination with PS (Naito et al., 2019; Sandhu et al., 2018). Since Ysp2-CT targets cortical ER without the Ysp2 PH^GRAM^ domain, we asked whether the Ysp2p PH^GRAM^ domain also binds sterol. From sequence conservation alone this seems likely, because the PH^GRAM^ domain is the most conserved region between GRAMD1B and Ysp2 (42% identity, 63% homology, compared to 22%/35% for the StARkin domain). Nevertheless, we tested whether the Ysp2 PH^GRAM^ domain is recruited by sterol.

We noted that the sequence flanking the PH^GRAM^ is highly electro-negative (Figure 3A). Previously, we had a broad definition of the PH^GRAM^ as a region of 211 residues (Gatta et al., 2015), but this region contains 19 net negative charges in disordered flanks of the folded domain (Figure 3A). Here, to avoid any construct being repelled by the anionic phospholipids of the PM, we cloned an electro-neutral region we called PH^GRAM^[short] (PH^GRAM^[S], 141 residues, 638-778, Figure 3A). When expressed in wild-type cells, this was diffusely present throughout the cytoplasm (Figure 3Bi). We wondered if one reason for the lack of PM targeting by PH^GRAM^[S] was that accessible free sterol is maintained at low levels in wild-type cells, which have the full complement of LAM proteins to remove free sterol. In human cells, ER-PM cortical targeting of GRAMD1B by PH^GRAM^ is self-limiting, because the StARkin domain transfers sterol away in a negative feedback loop (Sandhu et al., 2018). To see if the same occurs in yeast, we expressed PH^GRAM^[S] in the quadruple *lam1*Δ *ysp2*Δ *sip3*Δ *lam4*Δ strain (Sokolov et al., 2020). In these cells, PH^GRAM^[S] outlined the PM uniformly (Figure 3Bii). Since PH^GRAM^[S] binds not only sterol but also PS, we looked to see if cells lacking LAMs had altered PS distribution in the PM using the PS sensor Lactadherin C2 domain. This targeted the PM of wild-type cells and quadruple *lam* deleted cells similarly (Figure 3C) (Yeung et al., 2008). This indicates that the recruitment of PH^GRAM^[S] in the absence of LAMs (Figure 3Bii) most likely results from binding free sterol, as happens with human GRAMD1 (Naito et al., 2019; Sandhu et al., 2018). Confirmation will require demonstration that the purified domain binds directly to sterol in PS-rich liposomes.

**Figure 3:**
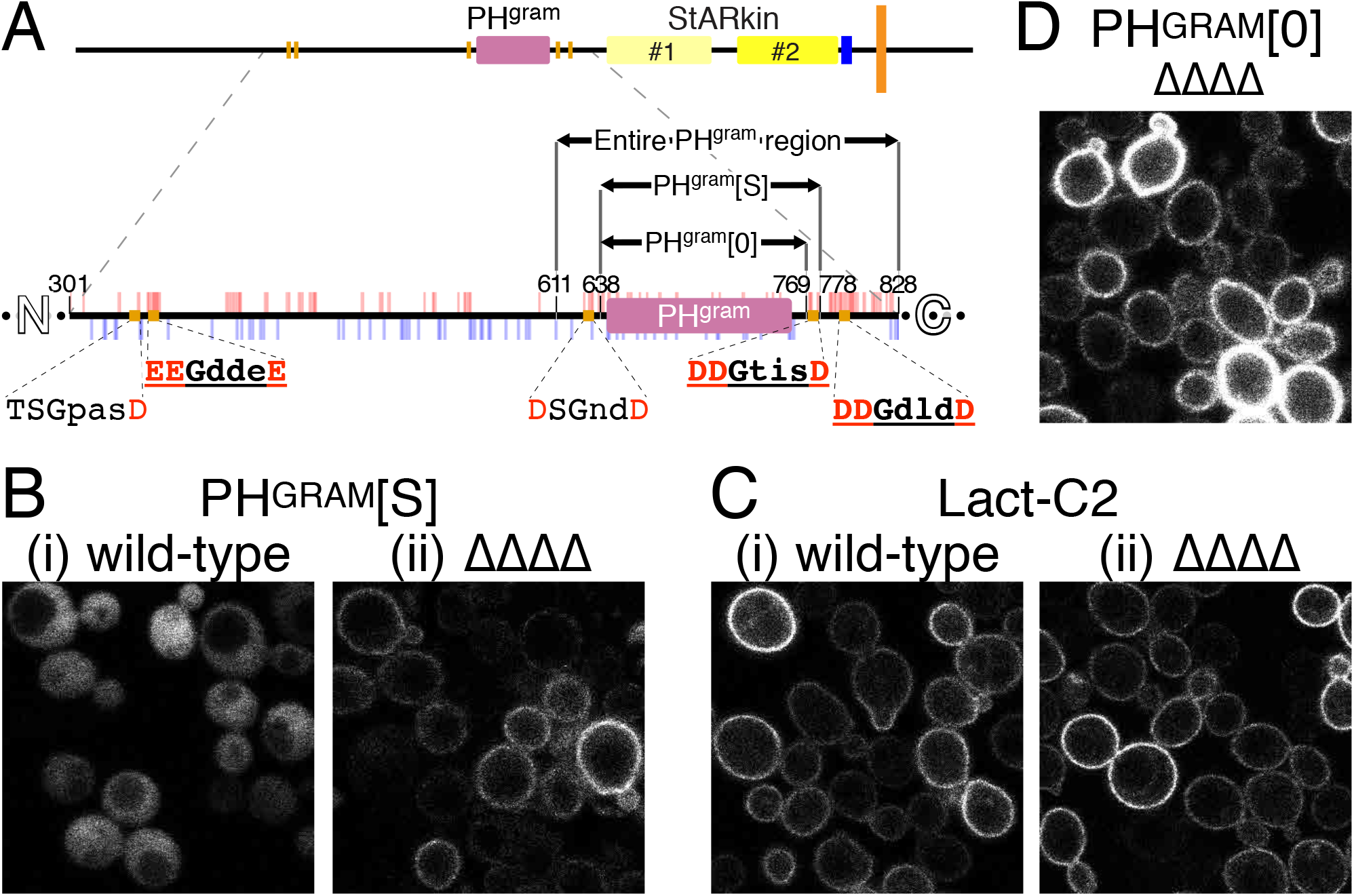
The PH^GRAM^ domain of Ysp2p is recruited to the PM when LAMs are absent and is surrounded by DSG degrons. **A**. Diagram of region surrounding PH^GRAM^ domain of Ysp2p showing 5 DSG degrons (orange rectangles) with close-up view (below) of residues 301-828. Also indicated are: degron sequences (bold/underlined è pre-activated); pattern of charges (acids and bases as red/blue ticks above/below); boundaries of PH^GRAM^ domain constructs: (a) entire PH^GRAM^ region (Gatta et al., 2015) that has 3 degrons and net 19 negative charges, (b) PH^GRAM^[S] that is electro-neutral and contains 1 degron, (c) PH^GRAM^[0] that contains no degrons (charge 3 positive). Other domains as in Figure 1. **B**. Short PH^GRAM^ domain construct (PH^GRAM^[S], containing residues 638-778) expressed in (i) wild-type cells and (ii) quadruple *lam1*Δ *ysp2*Δ *sip3*Δ*lam4*Δ deletes (ΔΔΔΔ). **C**. C2 domain from Lactadherin (Lact-C2) expressed in the same strains. **D.** As A(ii) but with PH^GRAM^[0].

Additionally, we noted that the negatively charged regions surrounding the PH^GRAM^ of Ysp2p contain three degrons for the highly conserved SCF-βTrCP ubiquitin ligase E3 ligase, with two other degrons further towards the N-terminus. The consensus for these degrons was originally defined as DSGxxS (Yaron et al., 1998) but has since been expanded (Bassermann and Pagano, 2010; Frescas and Pagano, 2008; Loveless et al., 2015), resulting in 64 variants (Supplementary Figure 2A). One of these degrons is present in PH^GRAM^[S] (^770^DDGTISD^776^, Figure 3A). To determine if this degron has an effect on construct stability, consistent with degron activity, we deleted it by truncating 9 residues from the C-terminus of PH^GRAM^[S] to create PH^GRAM^[0] with no degrons (Figure 3A). Expressed in quadruple *lam* delete cells PH^GRAM^[0] had a similar distribution as PH^GRAM^[S], but with higher levels of fluorescence per cell (Figure 3D, increase mean +58%, std. dev. ±8% n=3 experiments, 499 cells each on average). In wild-type cells, PH^GRAM^[0] was diffuse like PH^GRAM^[S] (not shown), and the levels were not increased (mean –8%, std. dev. ±2%, n=2 experiments, 375 cells each on average). These results suggest that SCF-βTrCP degrons surrounding PH^GRAM^ regulate the level of the construct when it is recruited to the PM, which may also apply to full-length Ysp2p protein, as with other degron-containing proteins (Low et al., 2014).

Ysp2 is not unique among LAMs in containing SCF-βTrCP degron motifs, as this motif is found in 4 yeast LAMs and in all human GRAMD1s (Supplementary Figure 2B). In addition, across the LAM family the most common degrons are of the form DDGxx(x)Z (where Z=D, E, S or T), being enriched 2-3 fold compared to eukaryotic proteins in general (Supplementary Figure 2C). Such substitution of S with phosphomimetic acidic D or E side-chains pre-activates the degrons (Kanemori et al., 2005). Indeed 3 of the motifs contained in Ysp2 are constitutively active, requiring no phosphorylation (Figure 3A). Although such motifs cannot be down-regulated by dephosphorylation, they can be regulated by protein-protein interactions (Jullien et al., 2011), which implies that protein interactions of the PH^GRAM^ region may regulate protein stability.

### 3. Fine mapping shows that the self-interaction interface in Ysp2-CT differs from GRAMD1B-CT 3A. The lumenal region of Ysp2 is necessary but not sufficient for punctate localisation

Next we set out to map which portions of Ysp2CT are responsible for its punctate distribution. As a negative control, we created a construct called here NegCT that contains the same overall skeleton as Ysp2CT, but with as many substitutions as possible to replace Ysp2 with generic sequence. To achieve this, we divided Ysp2CT into four regions: PBR, linker, TMH and lumenal region, and we produced generic versions of each region to serve as negative controls (Figure 2C, diagram; Figure 4, top). The negative control version of the PBR has 6 of the 10 positively charged Ysp2 residues substituted by negative charges. The generic version of the linker had 34 of 40 residues (85%) substituted with a portion of the ER-PM linker of Ist2p (Kralt et al., 2015), using 34 residues from Ist2 with no known protein interaction and no propensity to form an amphipathic helix (D’Ambrosio et al., 2020; Wong et al., 2021). For a generic TMH, the hydrophobic core of 16 residues was replaced with 16 leucines, which target the ER (Herzig et al., 2012; Munro, 1991). The lumenal region was simply truncated to leave 13 aa, losing >90% of its residues and all of its predicted secondary structural elements. We confirmed that NegCT acts as a negative control for punctate targeting by imaging GFP-NegCT, which showed low level linear fluorescence throughout the ER, peripheral and nuclear envelope without any punctate localisation (Figure 2C). There was strong intravacuolar accumulation of GFP, which could result from mis-sorting of the construct, resulting in exit from the ER and the secretory pathway to the vacuole, which is the default destination of the secretory pathway in yeast.

**Figure 4:**
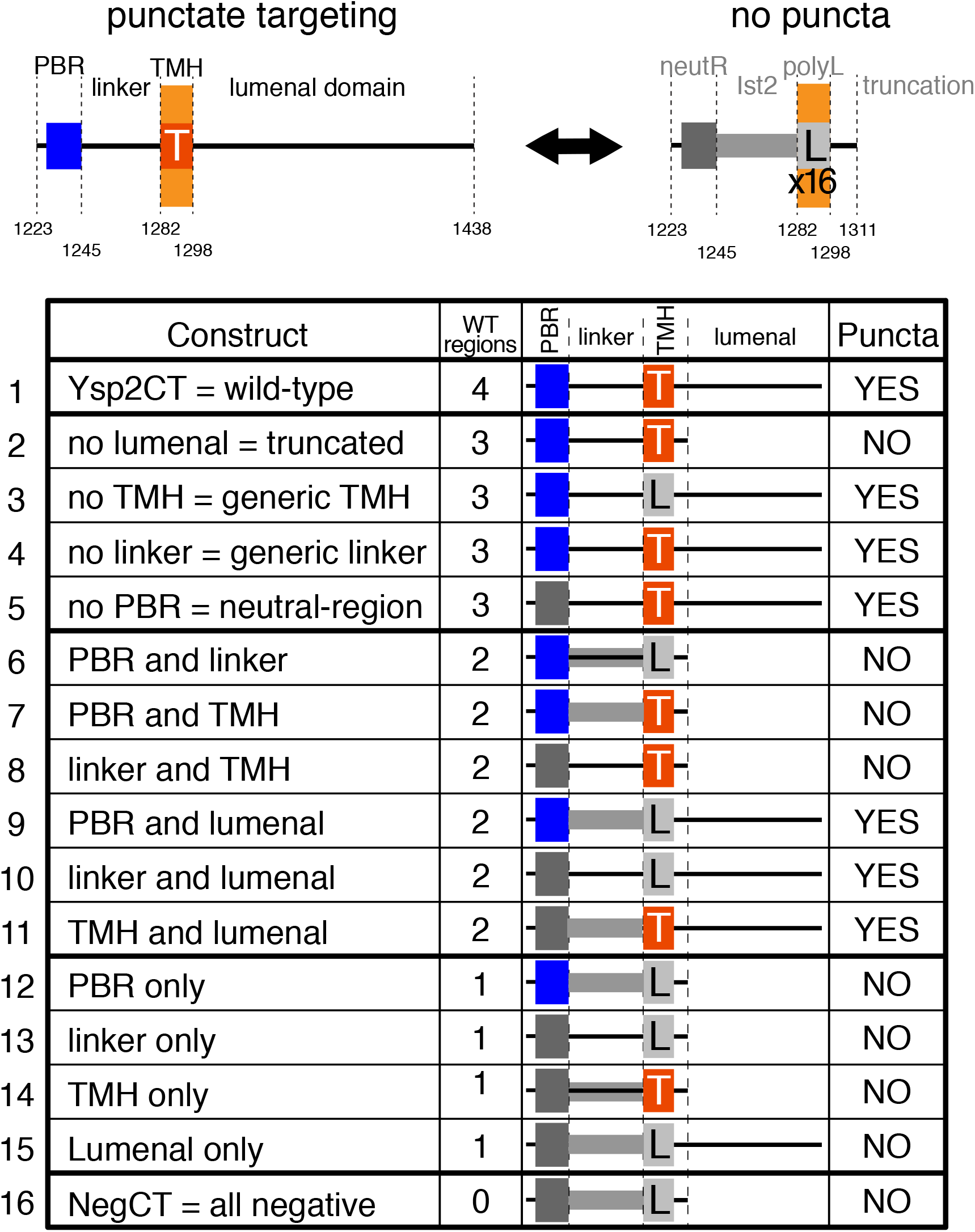
The lumenal region combined with any one of three other regions targets puncta. Summary of punctate targeting by 16 constructs made by all possible permutations of the four wild-type or the negative control regions. Top diagrams show the four regions in the wild-type construct (left) and their negative control counterparts. Targeting by these two constructs, with 4 and 0 of the wild-type (WT) regions, is shown in the top and bottom lines of the diagrammatic table below. Lines in-between show 14 other constructs with 3, 2 and 1 of the WT regions, illustrated by diagrams, and with presence of punctate targeting indicated.

We then used NegCT to test which regions in Ysp2CT mediate its punctate targeting. We swapped NegCT regions into the wild-type Ysp2CT construct one at a time (4 constructs), or in pairs (6 constructs) or with only a single region of the original Ysp2CT (4 constructs). The results are summarised in Figure 4. The key result was that the lumenal region is necessary for punctate targeting, in particular when this truncation was the only change (line 2). While necessary, the lumenal region was not sufficient for punctate targeting (line 15). Any other region of Ysp2CT (PBR, linker or TMH) restored punctate targeting when combined with the lumenal region (Figure 4, middle section). These results extends our previous finding that punctate targeting is inhibited by deleting most of the lumenal region (Gatta et al., 2015), and show that the adjacent portions of Ysp2CT are redundant in facilitating puncta formation.

### 3B. Punctate targeting of Ysp2 requires a lumenal sheet but not an amphipathic helix

We next examined punctate targeting of Ysp2p in the light of a previous study of human GRAMD1B, which identified a lumenal amphipathic helix as the key feature that mediates both homo-oligomerisation and hetero-oligomerisation with GRAMD1A (Naito et al., 2019). Substitutions in GRAMD1B that negate the amphipathicity of lumenal helix-2 (Figure 1C/D) reduced oligomer formation and prevented punctate targeting (Naito et al., 2019).

We first tested the amphipathic helix in the lumenal region of Ysp2CT (helix-1, Figure 1D, and Supplementary Figure 3A-C), with the result that deletion of this helix (together with helix-2) had no effect on punctate targeting (data not shown). The next Ysp2CT element we tested was helix-3 of the lumenal region, which is a strongly predicted coiled-coil (Figure 1D, Supplementary Figure 3D) (Wong and Levine, 2016), so might mediate oligomerisation. We introduced 3 levels of changes: 5 substitutions, a moderate deletion and also a major deletion (Supplementary Figure 3E/F). Rather than affecting punctate localisation, which was preserved, all three types of changes resulted in reduced expression, exemplified by the major deletion (Figure 5A). This indicates that the predicted coiled-coil is needed to stabilise Ysp2CT, but it is not necessary for puncta formation.

**Figure 5:**
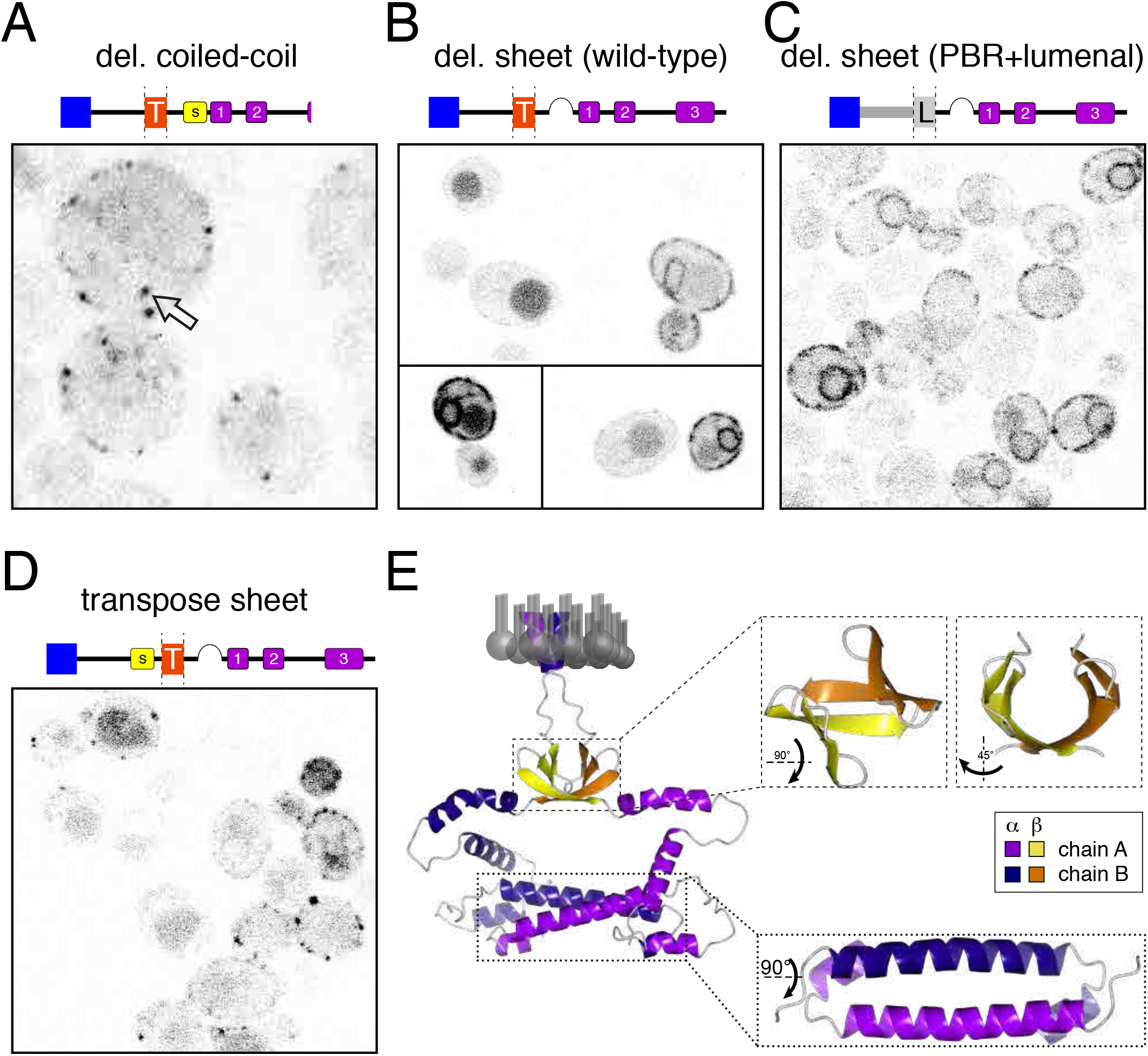
Punctate targeting by Ysp2CT requires a lumenal beta-sheet predicted to dimerise. **A.** Cells expressing GFP-tagged Ysp2CT with a major deletion of the lumenal amphipathic helix (helix-3) with a strong predicted propensity to form a coiled-coil (see Supplementary Figure 3A/F for details). Image is shown at 2x magnification compared to others to clarify the internal nature of some puncta (arrow). **B. and C.** The lumenal sheet in Ysp2CT was deleted, either as the sole substitution (B, compare to Figure 2A) or together with substitution of the linker and the TMH (C). Expression in B was only rarely detectable, so a gallery of three separate groups of expressing cells from different parts of the same image is shown. **D.** The sheet was transposed from its usual location in the lumenal region to the cytoplasmic linker, ending 6 residues before the TMH. The rest of the construct is as in B. Diagrams above images A–D summarise the construct expressed (see Figure 4 for key). Constructs were expressed in wild-type cells. **E.** Colabfold prediction of Ysp2CT dimer. Two dimerisation interfaces are shown in detail: the beta-sheet and helix 3, with rotation as indicated. α and ý structural elements in chains A and B are coloured as shown in the key; fog is used as a depth cue.

The only remaining element in the lumenal region is a region close to the TMH predicted to form a sheet of 3 β-strands (Figure 1C/D). Deletion of these led to uniform targeting across the ER with no detectable puncta, independently of whether other Ysp2CT regions were wild-type or generic (Figure 5B and C). To further test the role of the β-sheet, we transposed the 3 β-strands to the other side of the ER membrane, inserting them into cytoplasmic linker. This restored punctate targeting, though at reduced levels compared to wild-type Ysp2CT (Figure 5D). Overall, these results show that the major role of the lumenal region in creating the punctate localisation of Ysp2CT resides in the β-sheet.

We wondered if the β-sheet can oligomerise, and tested this *in silico* using ColabFold to predict a model from two copies of Ysp2CT (Mirdita et al., 2022). All of the five top-ranked models had two interfaces for dimerisation: (i) an antiparallel coiled-coil and (ii) a highly curved β sheet of 6 strands (Figure 5E).

Predicting structures for higher copy numbers of the sheet (n=3/4/6) only produced dimers with no higher order oligomerisation. Compared to the flat predicted sheet in the monomer (Figure 1D), the two sheets in the predicted dimer are more curved and form a globular domain of 42 residues (1310-1330 from each chain), dimensions ∼2×2×1.5 nm, with a largely hydrophobic core and subtle changes compared to the monomer, with more residues assigned to β-strands and greater strand curvature producing an almost spherical dimer. This structural prediction of a previously over-looked small all-β domain that can dimerise supports our results that self-interaction of Ysp2p involves the lumenal region, which has two interfaces.

### 3C. The TMH contributes a third self-interaction interface

Given that the lumenal domain is insufficient for punctate targeting, needing one of the other three regions we identified, we looked next for what those regions contribute. The previous study of targeting of GRAMD1B identified its TMH as contributing to self-interaction and punctate localisation (Naito et al., 2019). The simplest form of this self-interaction is dimerisation, which can be studied with structural bioinformatics. For example, the PREDDIMER server surveys all possible dimer conformations to identify the lowest energy conformation for any sequence (Polyansky et al., 2014). Applying PREDDIMER to the native LAM sequences showed that Ysp2p has a weaker propensity to dimerise than GRAMD1B, and both score much lower than a well-documented ER-located dimer such as VAPA, which contains a GxxxG motif, but still score well above background, which is represented by the generic TMH containing 16 leucines that we used in this study (Figure 6A) (Kim et al., 2010). To test if the TMH contributes to punctate targeting of Ysp2CT, we compared the effect of different substitutions on the localisation of the construct that depends on the TMH, accompanied by wild-type lumenal region with generic PBR and linker (Figure 6B). Punctate targeting, which was lost by adding the generic TMH (16 leucines, summarised Fig 4, line 14), was restored by adding known dimerisation motifs GxxxG, AxxxG to the generic TMH (Figure 6A) (Kirrbach et al., 2013). Since even AxxxG has a PREDIMER score marginally higher than that of Ysp2p, we used a construct containing a single glycine, (*i.e.*: motif equivalent to LxxxG) which has a PREDDIMER score lower than Ysp2p but similar to Lam4p. This too showed punctate targeting (Figure 6A and C). These results are consistent with self-interaction by the TMH in Ysp2CT contributing to puncta formation, similar to what has been shown for GRAMD1B, though direct evidence remains to be obtained.

**Figure 6:**
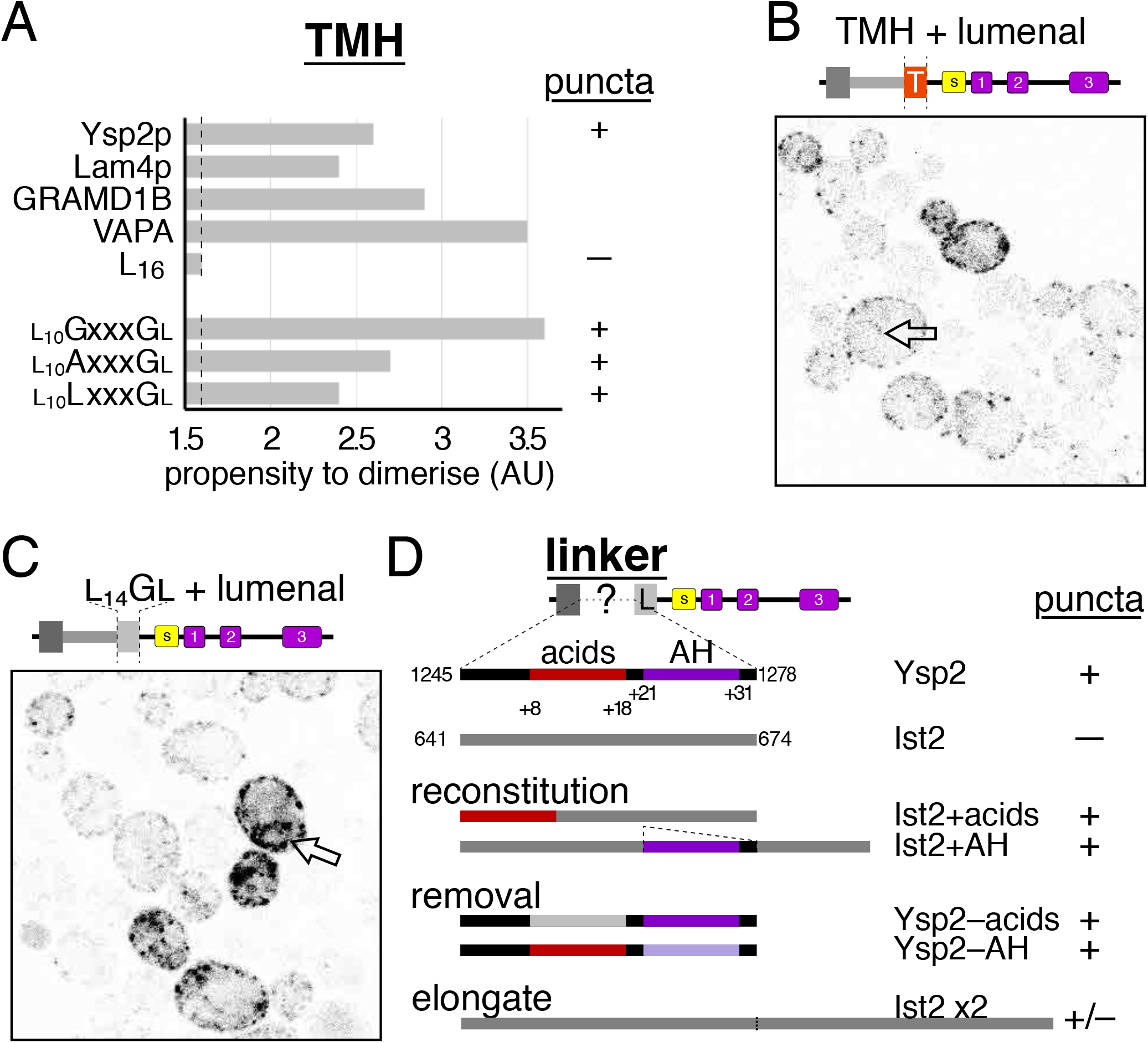
Punctate targeting of Ysp2CT is supported either by TMHs that dimerise or by longer linkers. **A**. Strength of dimerisation predicted by PREDDIMER for eight TMH sequences in LAMs (x3), VAP and generic constructs (x4), and corresponding observed formation of puncta in the five constructs tested in this study (“nd” means not done). PREDDIMER was used to report the maximum packing efficiency for putative TMHs, this relating to the propensity to dimerise (arbitrary units, AU) (Polyansky et al., 2012): Ysp2 1283-1302, Lam4 1204-1223, GRAMD1B 625-645 (21 residues), VAPA 226-245, L_16:_ 16 leucines followed by Ysp2 1299-1302 (RLLF). L_16_ scored 1.6, which is the same as 20 leucines (not shown). Therefore, this is taken to be the background signal (dashed line). L_16_ variants all have G at position 15, and have G, A or L at position 11. **B.** Construct with wild-type TMH and lumenal region. **C.** Construct in which only the lumenal region has Ysp2 sequence (rest is negative control) as in Figure 3C, but with a single substitution: leucine-15 of the TMHèglycine, which increases dimerisation propensity. Constructs with G or A at position 11 (producing GxxxG and AxxxG, with greater propensity to dimerise) also had punctate targeting (see part A). Arrows in B/C indicate fluorescence located on the nuclear envelope. **D.** Summary of localisation of 7 constructs that vary only in the linker region. The acidic region and the amphipathic helix (AH) in the Ysp2 linker were separately reconstituted into the Ist2 linker or they were removed from the Ysp2 linker. For details of sequence, key to colour scheme and images of cells expressing these constructs, see Supplementary Figure 4A–D. Note that the helix is weakly predicted by both PSI-PRED and AlphaFold. The final line summarises an elongated construct made by tandem repetition of the negative control linker was elongated (see Supplementary Figure 4E).

### 3D. Longer linker regions support punctate targeting better

Having found potential self-interaction interfaces in both the lumenal region and TMH, the linker and the PBR remain to be studied. Since inclusion of each of these alongside the lumenal region leads to punctate targeting (Figure 4), does that mean that they too can self-interact? How the PBR may work is addressed in the next section. Looking first at the linker, we noted two structural elements: an acidic stretch and an amphipathic helix (Figure 6D). We added each of these elements back into the negative control linker and we removed each of them from the wild-type linker (Supplementary Figure 4A). All four resulting constructs showed punctate targeting (summarised in Figure 6D, and see Supplementary Figure 4B/C). This indicates that neither the acidic stretch nor the amphipathic helix are the critical structural element that differentiates between the wild-type and the negative control linkers.

Although the wild-type and negative control sequences have the same number of residues, they may have different propensities to stretch/condense, resulting in different effective lengths. To test the importance of length, we duplicated 34 residues of the negative control linker in series, increasing it from 41 to 75 aa. The resulting construct localised to puncta (Supplementary Figure 4D). Punctate targeting initially increased by elongating the linker with sequential doublings of the linker to (x4/8 = 143/279 aa) (Supplementary Figure 4E), and then with even longer linkers (x16/32 = 551/1095 aa) punctate targeting plateaued. These results indicate that effective linker length is decisive for puncta formation.

### 4. ER-PM contacts containing Ysp2CT lack any structural specialisation

The results above suggest a molecular basis for the involvement of three of the four regions of Ysp2CT in its punctate localisation, but do not address the role of the PBR, which in other ER-PM contact site proteins is typically considered to interact with anionic phospholipids of the PM (Lees et al., 2017; Sohn et al., 2018). Ysp2p only has 40 residues between the TMH and PBR, indicating the PBR can at most reach 15 nm away from the ER (3.8 Å per residue for intrinsically disordered peptides (Pillardy et al., 2001)).

While an electron microscopic survey found that ER-PM distances were in general larger than 15 nm (West et al., 2011), tricalbins can induce highly curved peaks of cortical ER that come as close as 7-8 nm to the PM (Collado et al., 2019). To determine the morphology of contacts targeted by Ysp2, we used correlative light and electron microscopy (CLEM). We targeted GFP-Ysp2CT puncta to examine the associated ultrastructure of cortical ER (Figure 7A). ER at these sites was typical of cortical ER in general, consisting of both tubules and cisternae. We observed no enrichment of a specific ER architecture. Also, we did not observe cortical ER peaks, like those that have been associated with tricalbins (Collado et al., 2019). Electron tomograms of contact sites identified by the fluorescent signal of GFP-Ysp2CT were used to map the membrane planes of PM and ER in three dimensions across a region of ∼100-200 nm (Figure 7B). On average, the intermembrane distance was 23 nm (StdDev 7 nm, n=8 contact sites). This is similar to the average ER-PM distances that correlate with the presence of other proteins that bridge across ER-PM contacts, these being 20-22 nm (Hoffmann et al., 2019). Having measured the intermembrane distances in regular intervals, among the contact sites we also observed regions where the distance was less than 17.5 nm (Supplementary Figure 4).

**Figure 7:**
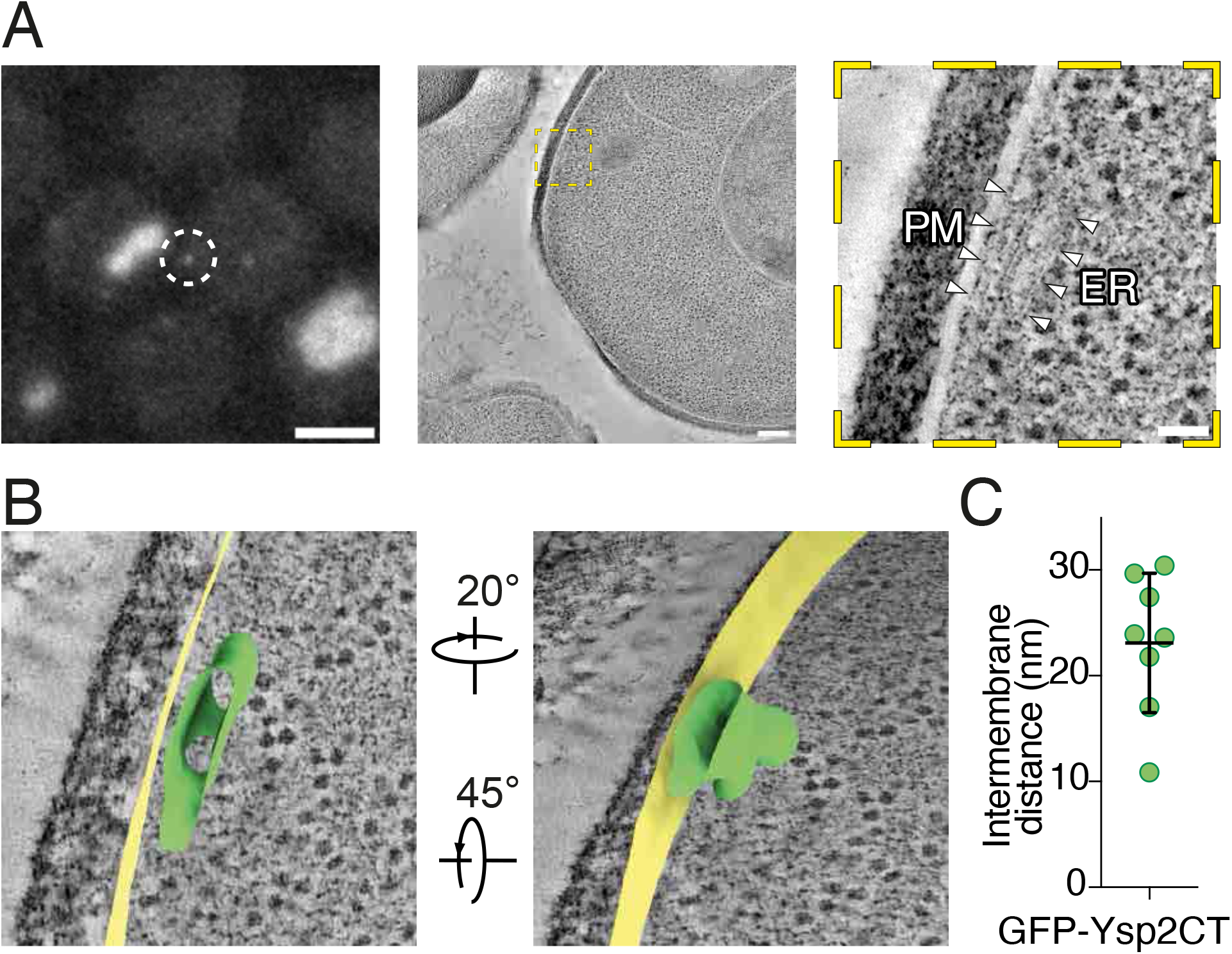
CLEM and ultrastructural analysis of ER-PM contact sites populated by GFP-tagged Ysp2CT. **A.** Workflow for CLEM. Left: fluorescence imaging, showing the average of fluorescence images from a z-stack acquired on resin sections of cells expressing Ysp2CT. A signal of interest is indicated by the dashed circle. Scale bar: 2 μm. Middle: A virtual slice through an electron tomogram acquired at the fluorescent signal indicated in the left image. Scale bar: 200 nm. Right: Zoomed-in view on the area corresponding to the localization of the fluorescent signal of interest, showing an ER-PM contact site. The ER membrane and the cytosolic leaflet of the PM are indicated by arrowheads. Scale bar: 50 nm. **B.** Segmentation model of the ER-PM contact site identified in A, showing the ultrastructure in 3D from two perspectives, related by the indicated rotations. PM is shown in yellow, ER in green. **C.** Intermembrane distances of 8 GFP-Ysp2-CT mediated contact sites that were identified by CLEM. Horizontal lines indicate mean and standard deviation.

Thus, Ysp2CT, while being highly concentrated in puncta, does not associate with a special ER structure, residing in contacts with an average ER-PM distance of 23 nm. We return to this in the Discussion, addressing whether there is enough variability within the ER-PM gap to allow the PBR to engage with anionic phospholipids in the PM.

### 5. Ysp2CT is required for stabilisation and punctate localisation of full-length Ysp2p

Knowing how regions in Ysp2CT determine its targeting, we looked to see what role these have for targeting the full length protein. *ysp2*Δ cells were transformed with GFP-tagged full-length Ysp2 constructs (Figure 8). Ysp2p with the wild-type C-terminus was entirely punctate and cortical, as expected (Ysp2^CT:WT^, Figure 8Ai). By comparison, the variant with all four C-terminal regions substituted by generic sequence (Ysp2^CT:generic^) was present at barely detectable levels, and localisation was not punctate (Figure 8Aii). Reintroduction of individual wild-type regions into Ysp2^CT:generic^ showed specific effects (Figure 8B). Adding back the PBR increased the signal considerably and produced a few puncta (Figure 8Bi). Including the linker region increased levels to a lesser extent than the PBR, and with fewer puncta (Figure 8Bii). Both other two regions failed to increase overall levels above that for Ysp2^CT:generic^, but they produced puncta, less bright for the TMH (Figure 8Biii) than for the lumenal region (Figure 8Biv). We next examined the effects of substituting individual regions of the C-terminus in Ysp2^CT:WT^ (Figure 8C). Neutralising the PBR had little or no effect (Figure 8Ci). Replacing the linker or the TMH with generic sequences led to increased expression without affecting punctate targeting (Figure 8Cii/iii). Uniquely, truncating the lumenal region changed the distribution to diffuse throughout the ER including the nuclear envelope, also increasing expression (Figure 8Civ). The punctate and diffuse ER distributions of the latter two constructs was confirmed in optical sections through the cell peripheries (Figure 8D).

**Figure 8:**
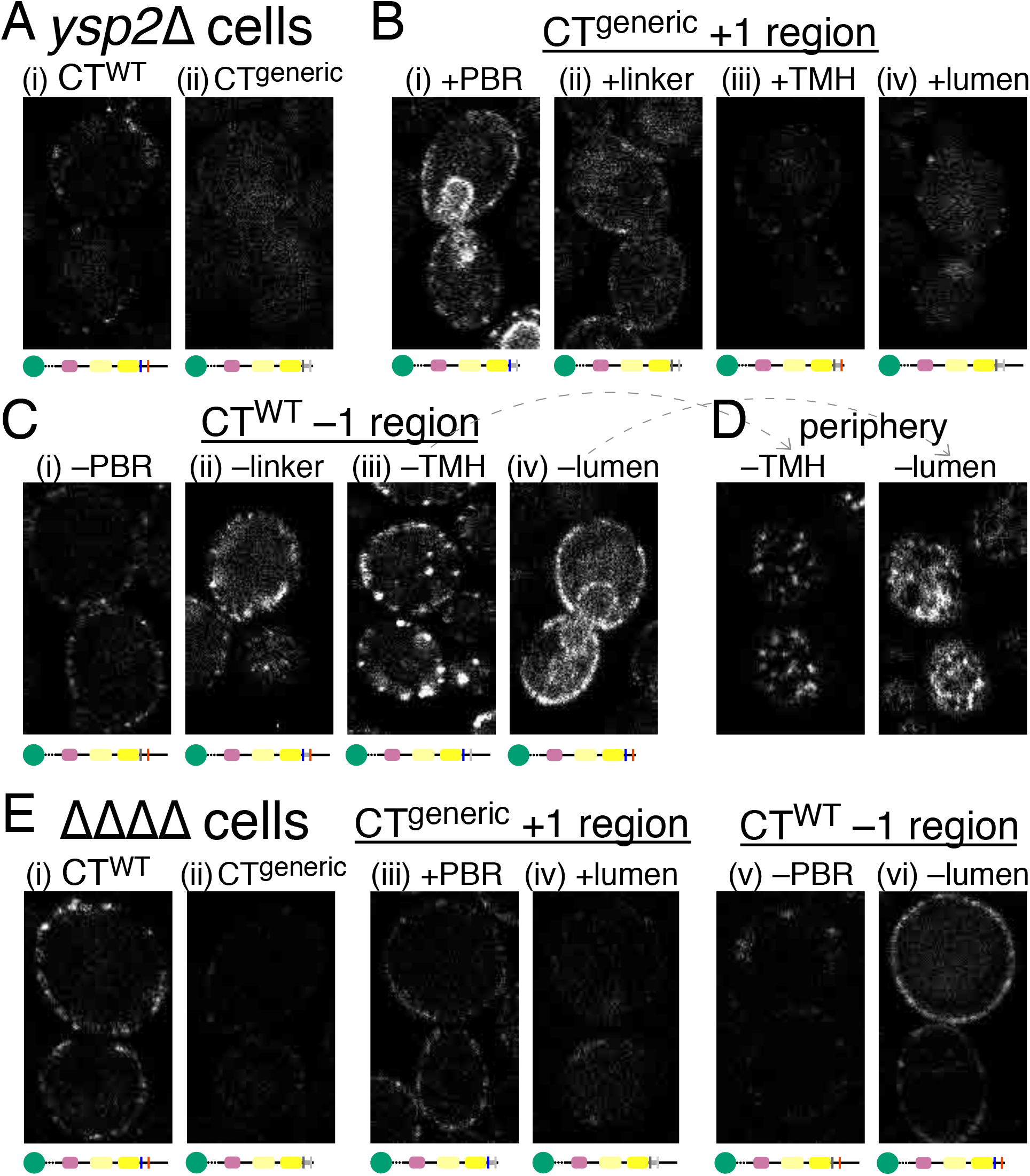
Full-length Ysp2p needs the lumenal region for punctate targeting and to support normal levels. **A and B**. Full-length GFP-tagged constructs expressed in *ysp2*Δ cells with varying Ysp2CT: A. CT either all wild-type (WT) or all generic, B. Generic C-terminus with single WT regions added. **C**. Wild-type C-terminus with single regions replaced by generic alternatives. **D**. peripheral sections of the constructs lacking TMH and lumenal regions from part C. In A(ii), Ysp2p with a fully generic CT formed no obvious puncta compared to the wild-type sequence (i).

A relevant question is whether these localisations of full-length Ysp2 constructs are affected by the other ER-PM LAMs, so we repeated the localisations in the quadruple null cells. These reproduced the major results seen above in *ysp2*Δ cells (Figure 8E). In general these results are consistent with what we found with the isolated C-terminus: the generic CT has low stability and does not form puncta; in any situation the lumenal region is a strong determinant of puncta formation while the PBR and linker are not. One clear difference for full length Ysp2 (compared to the isolated C-terminus) is that puncta were seen with either the TMH or the lumenal region alone, albeit with low expression levels (Figure 8Biii/iv). Such formation of puncta without the lumenal region, which was not seen for Ysp2CT alone, could be explained if an element within residues 1-1222 outside Ysp2CT enhances self-interaction.

### 6. Ysp2CT is required for function of full-length protein

We next looked at how the C-terminus affects the function of Ysp2p, assaying resistance of yeast to Amphotericin B (AmB), a polyene antifungal drug that kills cells through an initial interaction with free sterol in the PM (Anderson et al., 2014). Yeast resistance to AmB requires Ysp2p to transfer sterol away from the PM (Gatta et al., 2015; Murley et al., 2017; Topolska et al., 2020), and LAM function has also been correlated with AmB resistance more widely: in fission yeast (Marek et al., 2020), filamentous fungi (Zhu et al., 2021) and human cells (Ercan et al., 2021). Ysp2^CT:WT^ largely reconstituted the AmB resistance of *ysp2*Δ cells (Figure 9A, top 3 rows). In comparison, Ysp2^CT:generic^ was inactive, growth being the same as with an empty plasmid (Figure 9A, row 4). Next we added back each of the four CT regions individually to look for an increase in activity over Ysp2^CT:generic^ (Figure 9B). Re-introduction of the PBR significantly increased growth almost to the level of wild-type Ysp2p (Figure 9B, row 1). By comparison, adding either the linker or the TMH or the lumenal region had much lesser effects, with a clear increase in growth at a medium level of AmB (Figure 9B, rows 2–4, middle column, compare to Ysp2^CT:generic^), but little rescue at the high level of drug (right column).

**Figure 9:**
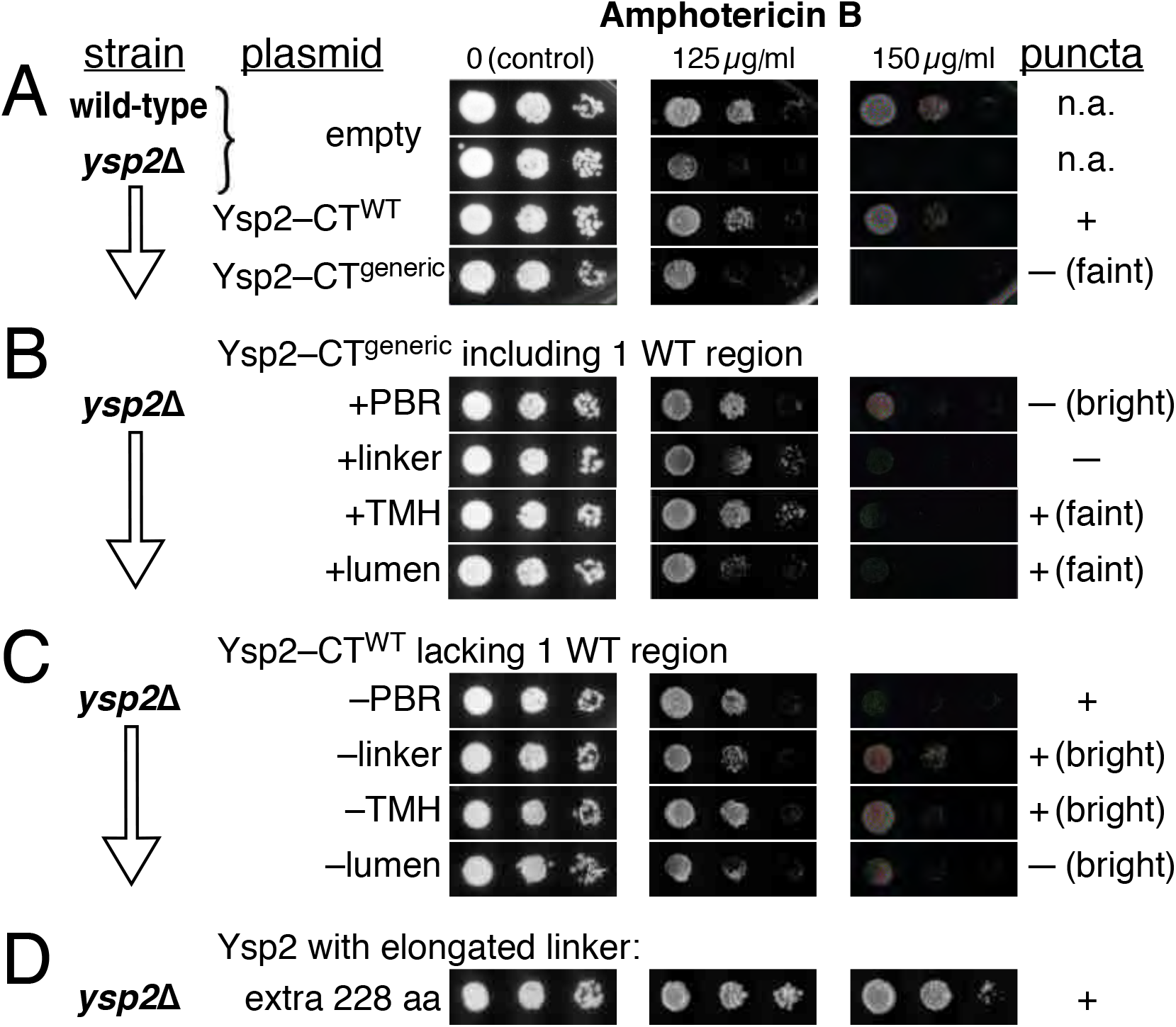
Full-length Ysp2p needs the wild-type C-terminus or specific regions within it to rescue AmB resistance. Resistance to Amphotericin B (AmB) rescued by full-length Ysp2p constructs with varying C-terminus. **A.** The wild type parent strain (line 1, only) is compared with *ysp2*Δ cells (all other lines) reconstituted with an empty plasmid (line 2), fully wild-type Ysp2p (WT, line 3) and Ysp2p with its C-terminus substitute with generic sequence (line 4). **B.** Ysp2p with generic C-terminus in *ysp2*Δ cells (as A, line 4) with single WT regions added. **C.** Ysp2p with WT C-terminus in *ysp2*Δ cells (as A, line 3) with single regions replaced by generic alternatives. Concentrations of AmB that partially and completely inhibited growth with an empty plasmid were 125 µg/ml (medium) and 150 µg/ml (high) respectively. (iv) *ysp2*Δ cells expressing Ysp2p with C-terminus containing 8 copies of generic linker, with one other region generic: the TMH. The amplified region of 34 aa adds an extra 238 aa to the linker (normally 41 aa, now 279 aa). All constructs are tagged with GFP+myc and expressed under the *YSP2* promoter (see Supplementary Table 1). White lines indicate where different parts of images from the same or parallel plates have been re-arranged.

We also took the approach of replacing single regions in Ysp2^CT:WT^ to see if any are essential for function (Figure 9C). Neutralising the PBR had the strongest effect, as without the cationic charge there was no growth at high AmB (Figure 9C, row 1). In contrast, the wild-type linker and TMH were dispensable (Figure 9C, rows 2, 3). Truncating the lumenal region caused a partial growth defect, most clearly seen at moderate AmB levels (Figure 9C, row 4, middle column). A further set of variants we tested were CT constructs with elongated linkers (Supplementary Figure 4A/E). Longer linkers (x2/4/8 copies of generic linker) led to a gain of function above that seen for wild-type Ysp2p (for example, 8 copies: Figure 9D).

These results indicate that the C-terminal region is necessary for function of Ysp2p, and that within the C-terminal region the critical portion for function is the PBR. The most obvious correlation between function of the different constructs (Figure 9) and their appearance by confocal microscopy (Figure 8) is that function positively correlates with level of expression, likely caused by stabilisation of the construct. More detailed correlations will be discussed below.

## Discussion

Here we carried out an in depth analysis of our previous finding that the C-terminus of Ysp2p localises to cortical ER puncta similar to the full-length protein (Gatta et al., 2015), which parallels work on targeting by the C-terminus of GRAMD1B in human cells (Naito et al., 2019). Ysp2 C-terminal determinants can now be ascribed specific roles relating to biophysical properties such as dimerisation, length and charge. Importantly, punctate targeting of Ysp2CT does not require partner proteins, in particular other LAMs (Lam1p/Sip3/Lam4p) with which it forms complexes. Although Ysp2p also forms complexes with Laf1p/Dgr2p (Murley et al., 2017), this pair of cytoplasmic proteins are recruited by LAMs not *vice versa* (Topolska et al., 2020). Thus, Ysp2 targeting is largely self-organising. Our approach to identify the active components within Ysp2CT was to substitute it with generic counterparts. The 100% generic C-terminus was wholly negative in terms of both punctate localisation and supporting protein stabilisation/function. We cannot exclude that this results from a toxic, dominant negative effect of the generic sequences we chose. However, when each of the four regions were examined on their own and in pairs, none were toxic, so the approach allowed approximate functions to be attributed to each region in Ysp2CT.

For punctate targeting by Ysp2CT, the lumenal region is the only part that is necessary, though it is not sufficient, so it is not a targeting sequence in the strict sense. The lumenal region of Ysp2CT contains two potential oligomerisation interfaces that could be responsible for accumulation in puncta. One of these is helical as in GRAMD1B (Naito et al., 2019). However, the other is the critical portion for punctate localisation, and this is a β-sheet (21 residues making 3 strands in each monomer) that is predicted by Alphafold multimer to be able to dimerise, so we suggest that this is a dimerisaton interface. This newly described small domain is also present in Lam4p and in LAMs in other Saccharomycetes such as *Pachysolen tannophilus*, separated from *S. cerevisiae* by ∼200 Myr, and more diverse non-Ascomycota species, such as *Ustilago maydis* (∼600 Myr separated), however it is far from universal, being absent from *Chaetomium* and *S. pombe*. The TMH in Ycp2CT is another dimerising region that contributes to punctate targeting (Figure 6), as it does in GRAMD1B (Naito et al., 2019). The precise residues that mediate dimerisation in Ysp2p and GRAMD1B have not been tested, but both TMHs share the short sequence S-xxx-C, which is a variant of the small-xxx-small dimerisation motif (Chen and Chou, 2017).

How the TMH and lumenal regions combine to produce punctate targeting is not clear. The three dimerisation interfaces (sheet, coiled-coil, TMH) might theoretically all bind the same partner (A=B), or they could be arranged to produce a chain (–A=B–C=D– *etc*.), or along with chains some proteins may interact with 3 different partners (A–B**^/^**^C^—D), which would produce a cluster.

The two regions that come before the TMH, the PBR and linker, also have an impact on Ysp2CT localisation and Ysp2 function. These two regions together consist of 60 residues (Figure 1C, Supplementary Figure 5A). At 0.38 nm per peptide bond (Pillardy et al., 2001), they allow a maximum stretch of 23 nm from ER to PM for the StARkin domain to acquire PM sterol (Supplementary Figure 5B, left). However, if the PBR fully engages with phospholipids on the PM, then only 40 residues would be available to reach from ER to PM (maximum distance 15 nm) (Supplementary Figure 5B, right). We also envisage an intermediate configuration, where the 9 PBR residues nearest to the StARkin domain (7 lysine/arginines) interact with the PM, leaving 46 aa to stretch across the gap (maximum 17.5 nm) (Supplementary Figure 5B, middle). CLEM of the Ysp2CT-rich puncta showed that these sites have the structure typical of cortical ER, with an average intermembrane distance of 23 nm (Figure 7). The lack of distinctive features is interesting given that LAM proteins are located to ER-PM contacts that are distinct from other ER-PM bridging proteins both in yeast (Quon et al., 2018) and in mammalian cells (Besprozvannaya et al., 2018).

How might the key domains of Ysp2p be arranged given the structure of the contacts? We speculate that the PBR within Ysp2CT, at least in part, engages with anionic PM lipids at regions where the ER membrane undulates so that the ER-PM gap is ≤17.5 nm, which are found in every contact (Supplementary Figure 5C). For full length Ysp2p, the intrinsically disordered peptide at its N-terminus could limit the number of PBRs within each punctum through its large hydrodynamic radius, which prevents crowding (Jamecna and Antonny, 2021). Consistent with this, elongation of linkers would allow PM engagement for more PBRs in the same cluster, which might underlie our finding that artificial elongation of the linker creates larger complexes (Supplementary Figure 4D/E).

Looking at full length Ysp2 for roles of C-terminal regions, the clearest correlation was between level of expression and function (Figures 8 and 9). Further examining if function is linked to punctate localisation, we found that only the PBR could restore function as a single WT region added back to Ysp2^CT:generic^, despite Ysp2^CT:generic+PBR^ being diffuse within the ER. Conversely, both Ysp2^CT:generic+TMH^ and Ysp2^CT:generic+lumen^ were barely functional even though they had similar punctate distributions to Ysp2^CT:WT^. Removing single regions from Ysp2^CT:WT^ again showed that the PBR was most critical for function, while in contrast removing the lumenal domain had little functional impact, possibly because of the high levels of Ysp2^CT:WT–lumen^, even though it was not punctate. Thus, despite the punctate localisation of Ysp2CT having multiple determinants and being a conserved feature of LAMs (Naito et al., 2019), it does not positively correlate with function.

Finally, we can attempt to link results regarding Ysp2CT with the other topic we addressed, namely the targeting of the PH^GRAM^ domain. Expressed on its own the PH^GRAM^ domain of Ysp2p targets the PM in LAM-deleted cells, consistent with it binding sterol as does the PH^GRAM^ of GRAMD1B (Naito et al., 2019; Sandhu et al., 2018), which is unsurprising since this domain is relatively well conserved. In contrast to GRAMD1B, which needs its PH^GRAM^ to translocate to the cell cortex (Naito et al., 2019; Sandhu et al., 2018), in Ysp2p the C-terminus targets this site constitutively, indicating an alternative function for Ysp2-PH^GRAM^. The degrons flanking PH^GRAM^ suggest that this function might be in negative regulation of Ysp2p, initiate degradation through recruiting the SCF-βTrCP ubiquitin ligase (in yeast: Met30p). We speculate that this function of PH^GRAM^ domain is itself linked to punctate targeting, in that both are required for negative regulation of Ysp2p. This is consistent with our finding that punctate targeting does not positively correlate with function. Instead, the most obvious correlation for the full-length protein is that it requires the PBR for high level expression (Figure 8B/C) and for function (Figure 9B/C), but the PBR negatively correlates with punctate targeting (Figure 8B/C), indicating that punctate targeting is overall inhibitory. Here, when we elongated the linker between the StARkin domain and the TMH the full length protein became more active than the wild-type (Figure 9A/D), which agrees with our previous observations for anchored constructs with the StARkin domain only (Gatta et al., 2015). We speculate that wild-type LAM/GRAMD proteins are under evolutionary pressure to anchor their active lipid transfer domains in puncta close to the ER membrane, imposing short cytoplasmic linkers (Wong and Levine, 2016), so they can be negatively regulated.

## Methods

### Strains

*ysp2*Δ is from the Euroscarf delete collection in BY4741 background (the wild-type parent). Quadruple delete strain: *lam1*Δ *ysp2*Δ *sip3*Δ *lam4*Δ strain in W303 background is as published (Sokolov et al., 2020). We verified the deletion of all four LAMs by genomic PCR of their LAM domains (data not shown). For cryo-EM, the W303 yeast strain was used.

### Constructs

**<u>Ysp2CT constructs</u>**: Ysp2 1223-1438, with a single change: arginine inserted between S1244 and P1245, were expressed in pRS416 with the *YSP2* promoter (510 bp before ATG), then an open-reading frame containing (i) monomeric Umikinoko Green fluorescent protein (mUkG) (Tsutsui et al., 2008), codon-optimised for expression in *S. cerevisiae* (Kaishima et al., 2016) and hereafter abbreviated to GFP, (ii) myc tag with short linkers (vtklgs<u>EQKLISEEDL</u>sn), (iii)-(vi) 4 regions of CT^WT^ or CT^generic^ (PBR-linker-TMH-lumenal) joined in series (see below for sequences and details of joining). There was no terminator. Ysp2CT^WT^ contained 89 silent changes to increase its Codon Adaptation Index (CAI) from 0.135 (wild-type) to 0.503. Large DNA fragments were purchased from IDT, and oligonucleotides from Sigma-Aldrich. All parts of constructs derived from purchased DNA products were verified by sequencing.

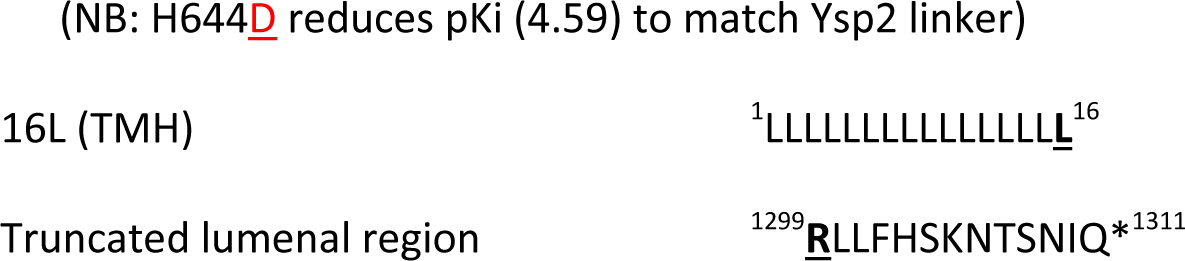
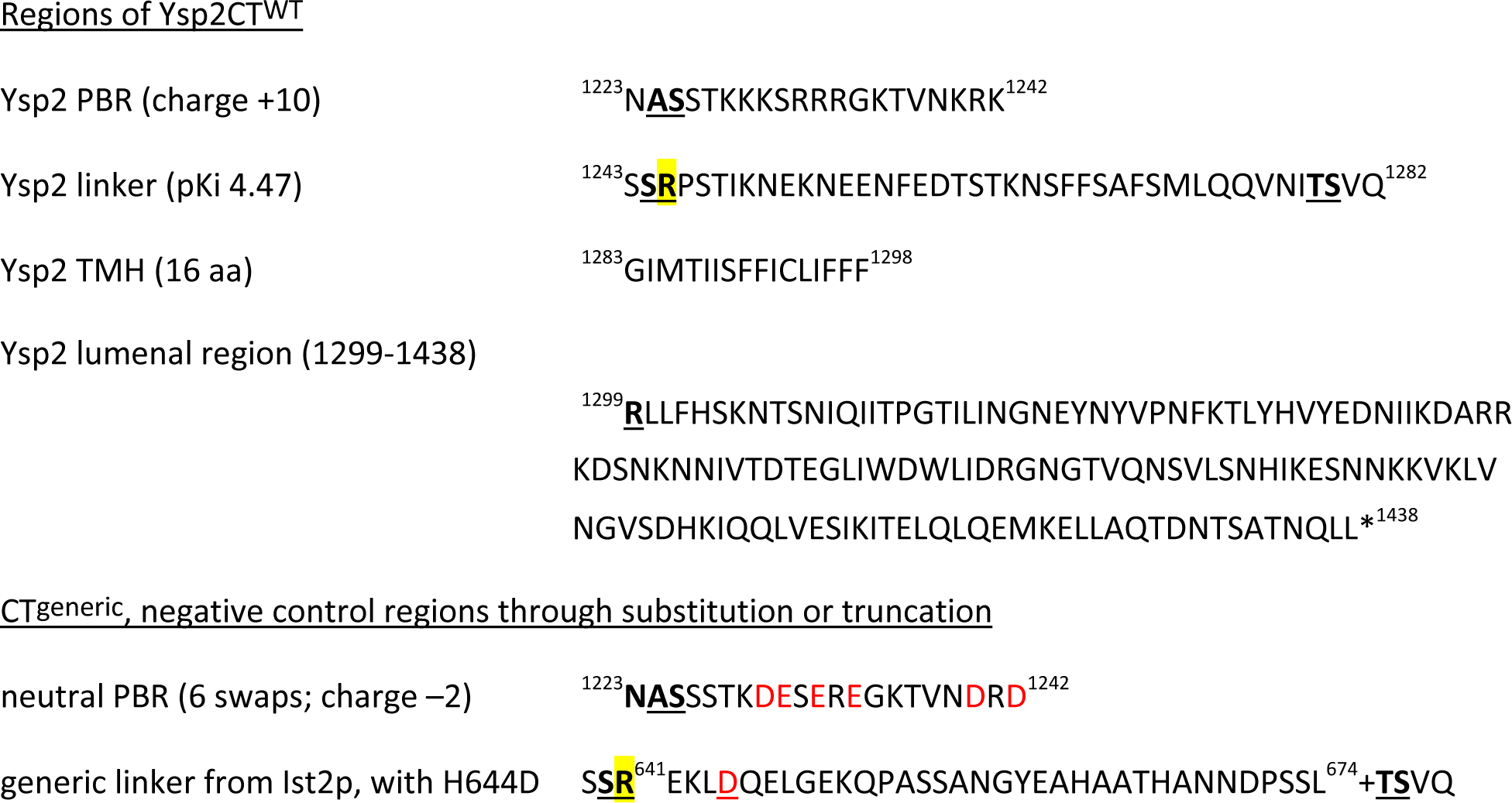

<u>Joining regions</u>: 16 constructs were made by combining Ysp2 WT regions with negative control regions together in all possible combinations: (NheI) – PBR – (XbaI) – linker – (SpeI) – TMH – (Bsu36I) – lumenal region. The restriction enzymes in brackets code for bold/underlined residues at regional joins, which are native to the Ysp2p sequence except for the added arginine (in yellow): NheI: ^1224^**<u>AS</u>**^1225^; XbaI: ^1244^**<u>S</u>** + added **R**; SpeI: ^1279^**<u>TS</u>**^1280^; Bsu36I: ^1298^**<u>LR</u>**^1299^.

**<u>Full length constructs</u>**: Ysp2 residues 1-1222 were added between GFP-myc and the Nhe1 site of Ysp2CT constructs.

#### Other constructs

Lactadherin-C2-GFP was expressed using the GPD promoter in pRS416, plasmid acquired from Addgene (Yeung et al., 2008). mUkG was integrated downstream of Cse4p in a construct that contains (i) the final 366 bp of the *CSE4* open-reading frame followed 24 bp encoding an 8 aa linker of SSGSSGSS, and then mUkG-myc (as in constructs above); (ii) the selection marker *Candida URA3* gene; and (ii) the *CSE4* 3’UTR (283 bp), this being adapted from an integration construct we used previously (Castro et al., 2022).

### Imaging

#### Microscopy

Yeast growing in mid-log phase were pelleted by centrifugation and mounted between slides and coverslips for microscopy on a Leica SP-8 confocal microscope using the 488 nm excitation line with varying laser power and pinhole of 1-1.5 Airey Units.

#### Image Treatment

Images were treated only by changing brightness and background, using the “levels” function in Adobe Photoshop to change the bottom and top value of 256 channels, with gamma kept at 1.0. For confocal images of cells expressing GFP-tagged constructs, identical re-scaling was used within each individual experiment. In some cases, regions were cropped after rotation of the image (15/30/45°). Inclusion of cells from multiple regions of an original image is indicated by internal white lines (for example, Figure 5B).

To assess brightness of green fluorescence per cell, whole random fields of cells were analysed for total signal after subtraction of background, and cells counted (average 410 (std. dev. 52) per condition) to arrive at signal per cell (arbitrary units). Ratios between strains were measured in three independent experiments. For scanned images of cells spotted onto agar, black background was exactly the same within each experiment but marginally different between experiments.

#### Determining copy number of Ysp2CT per punctum

Single confocal sections of UkG-Ysp2CT and Cse4-mUkG imaged under identical settings on the same day were used to avoid bleaching. Regions of interest containing a single punctum from >100 cells was assessed in NIH Image to obtain their fluorescence above background.

### Predicting structure

Colabfold (Mirdita et al., 2022) was used online at <u>colab.research.google.com/github/sokrypton/ColabFold/blob/main/AlphaFold2.ipynb</u>, submitting 382 aa from the C-terminus of Ysp2 including the second StARkin domain (1057-1438), with refinement of structures with Amber-Relax relaxation turned on. Subsequently, predictions were made for two or more chains, omitting the PBR and linker.

### Net charge of protein regions

this was determined from the amino acid count as (D+E)-(K+R+H), which assumes all histidines are positively charged at the neutral pH of the cytoplasm.

### Correlative light and electron microscopy (CLEM) and analysis of resulting electron tomograms

Ysp2CT-GFP under the YSP2 promoter was expressed by integrating plasmid pRS405 at LEU2 in the W303 strain. The colony selected for liquid culture had bright, uniform expression, indicating integration of more than one copy. The culture was grown in selective minimal medium at 25°C to mid-log phase and prepared for CLEM as described before (Kukulski et al., 2012), pelleted by vacuum filtration and high-pressure frozen with a Leica EMPACT-2. The The resulting samples were then freeze substituted in acetone with 0.05% (w/v) uranyl acetate, first by gently shaking on dry ice for 3 hrs (McDonald and Webb, 2011) before being placed into the Leica EM AFS2 at −90°C for 17 hrs. The temperature was subsequently raised to −45°C (7.5°C/hr) before washing with acetone three times. This was followed by gradual infiltration with Lowicryl HM20 (Polysciences) diluted in acetone at increasing concentrations (10%, 25%, 50%, 75%, 4 hrs each), while raising the temperature to −25°C. After that, three exchanges of 100% Lowicryl 4 hrs each were carried out and the resin polymerized by UV at −25°C for 48 hrs. The temperature was brought up to 20°C (5°C/hr) and UV polymerization continued for 48 hrs.

The resin-embedded samples were cut on a Reichert-Jung Ultracut E ultramicrotome equipped with a DiATOME ultra 45° diamond knife to produce 200 nm thin sections that were applied to carbon-coated 200 mesh copper EM grids (Agar Scientific). TetraSpeck 0.1 μm microspheres (Thermo Fisher Scientific) diluted in PBS were adsorbed onto the grid. For fluorescence imaging, grids were sandwiched between a coverslip and glass slide with a drop of PBS. Green channel z-stacks of the sections were acquired on a Nikon Eclipse Ti2 microscope using a CFI Apochromat TIRF 100×/1.49 NA oil objective and a Photometrics Prime BSI sCMOS camera. A Lumencor SpectraX light source (Chroma) with a 470 nm LED was used for excitation through a quad band filter set 89000 ET Sedat Quad (Chroma) including ET490/20x, 89100bs, ET525/36m. An emission filter wheel (Nikon) was set to 535 nm.

For electron tomography, 15 nm protein A-coupled gold beads were applied to both sides of the grid as fiducial markers for reconstruction and the sample was post-stained with Reynolds lead citrate. Grids were placed in a high-tilt holder (Model 2020, Fischione) which was inserted into a FEI Tecnai Spirit TEM operated at 80 kV with a Tungsten filament. Dual axis tilt series were recorded on a FEI 4k Eagle camera over a tilt range from −55° to 55° with 1° increment at a pixel size of 1.2 nm using SerialEM (Mastronarde, 2005). For correlation to fluorescence images, lower magnification montages at 3.3 nm pixel size were collected. Tomograms were reconstructed using IMOD (Kremer et al., 1996). TetraSpeck-based correlation was performed using MATLAB-based scripts described in (Kukulski et al., 2011).

ER-PM contact sites marked by the GFP-Ysp2CT signal were parametrized by manually defining points on both the ER and the PM approximately every 6 nm along the z axis in the tomographic volume. Using MATLAB scripts described in (Ganeva et al., 2023), surfaces corresponding to ER and PM bilayers were rendered through these points and the distances between them measured every 5 nm. The extracted distances were then analyzed with the R software package (version 4.0.4) using gglpot2 (version 3.3.3) and ggpubr (version 0.4.0). A probability density function was calculated from all measured distances at each contact site and the peak value taken as the final measurement for that contact site. In total, 8 contact sites were analyzed and are presented in a scatter plot generated in GraphPad Prism (version 9.4.1). Segmentation models of the ER and PM shown in Figure 7 were manually produced using IMOD (Kremer et al., 1996) and rendered in UCSF ChimeraX (Pettersen et al., 2021).

### Amphotericin B (AmB) resistance assay

15-fold dilutions of cells from up to 16 strains were spotted onto 9 cm plates, one with no AmB and others containing different concentrations between 50 µg/ml and 250 µg/ml added to medium cooled to below 50 °C from a 250 mg/ml stock in water (Sigma-Aldrich) aliquoted to reduce freeze-thaw cycles and stored at -20°C. To identify AmB concentrations where resistance was restored, between 2 and 4 concentrations were chosen for each 2-fold increase in AmB. After 48 hours growth at 30°C plates were scanned (≥300 dpi, ≥105 pixels per spot). Images shown are representative of three independent repeats.

### Identifying Degrons

6693 full length sequences with the LAM/Aster/VASt domains were downloaded from Pfam (Pf16016) and reduced in redundancy to 2244 sequences, using MMseqs2 with default settings (Mirdita et al., 2019), and supplemented by all human and yeast LAM sequences, making 2253 sequences (1926163 residues, mean 855 residues/protein). SCANPROSITE found 4641 hits in 1714 sequences for the string [DEST]-[DEST]-G-x(2,3)-[DEST]. As a control for eukaryotic proteins in general we started with the entire human genome of 20613 sequences downloaded from Uniprot (11406376 residues, 553 residues/protein), which led to 22855 hits in 10000 non-redundant sequences. Hits were then parsed for their composition in first, second and final positions. The number of hits in each of the 64 possible variants were expressed as fold excess of hits per residue in LAMs compared to hits per residue in the human proteome.

## Acknowledgements

We thank Svyatoslav Sokolov and Anant Menon for providing the *lam1*Δ *ysp2*Δ *sip3*Δ *lam4*Δ strain. G A-B was funded by the BBSRC (Grant BB/M011801/1). Work in the lab of W.K. was supported by the Swiss National Science Foundation (project 201158) and the University of Bern.

## Supplementary Figures

**Supplementary Figure 1:**
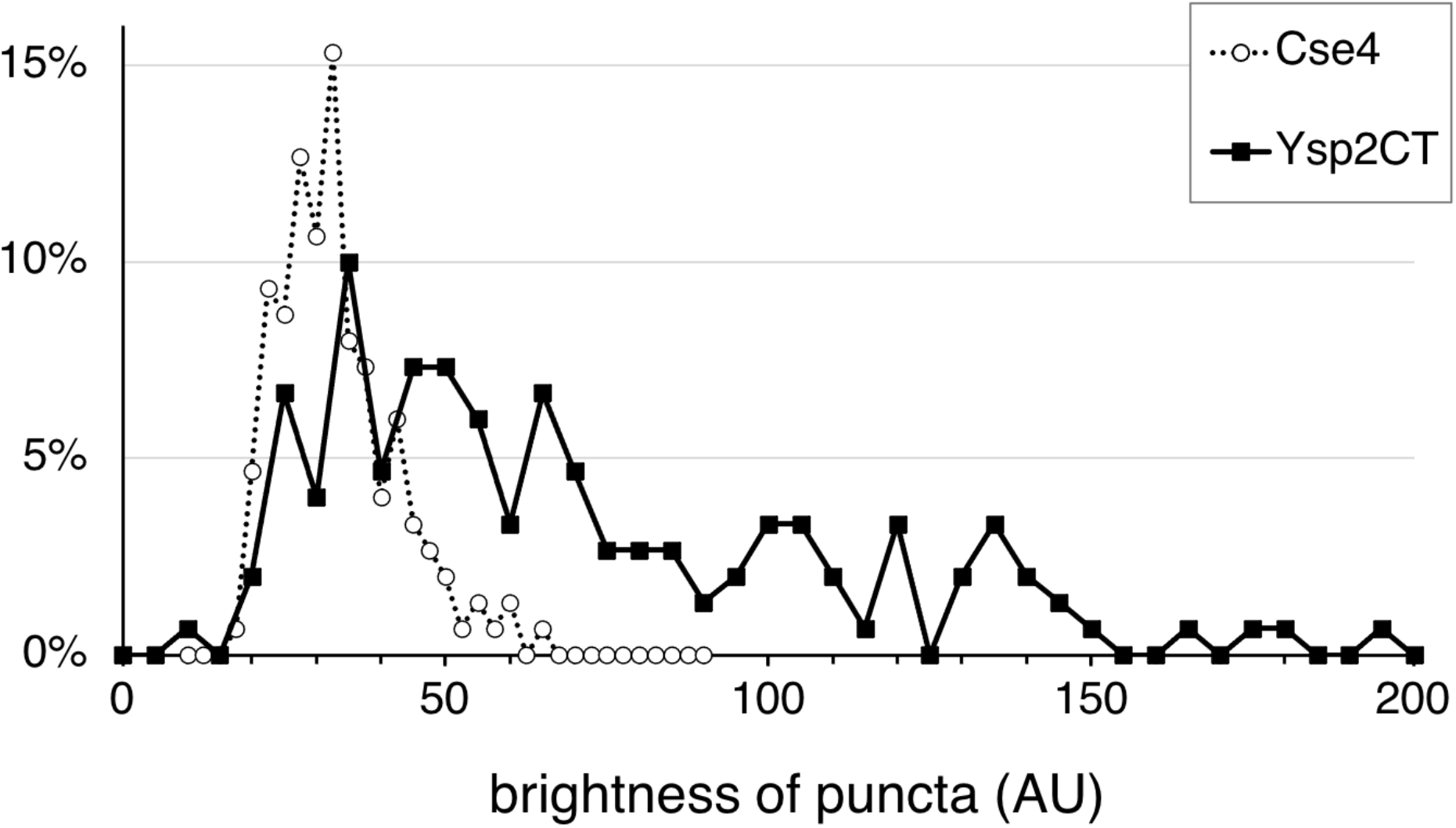
Brightness of puncta containing Cse4 and Ysp2CT. Fluorescence in puncta expressing either Cse4-mUkG at the genomic locus or mUkG-YspCT (wild-type) was quantified. n>100 in both sets.

**Supplementary Figure 2:**
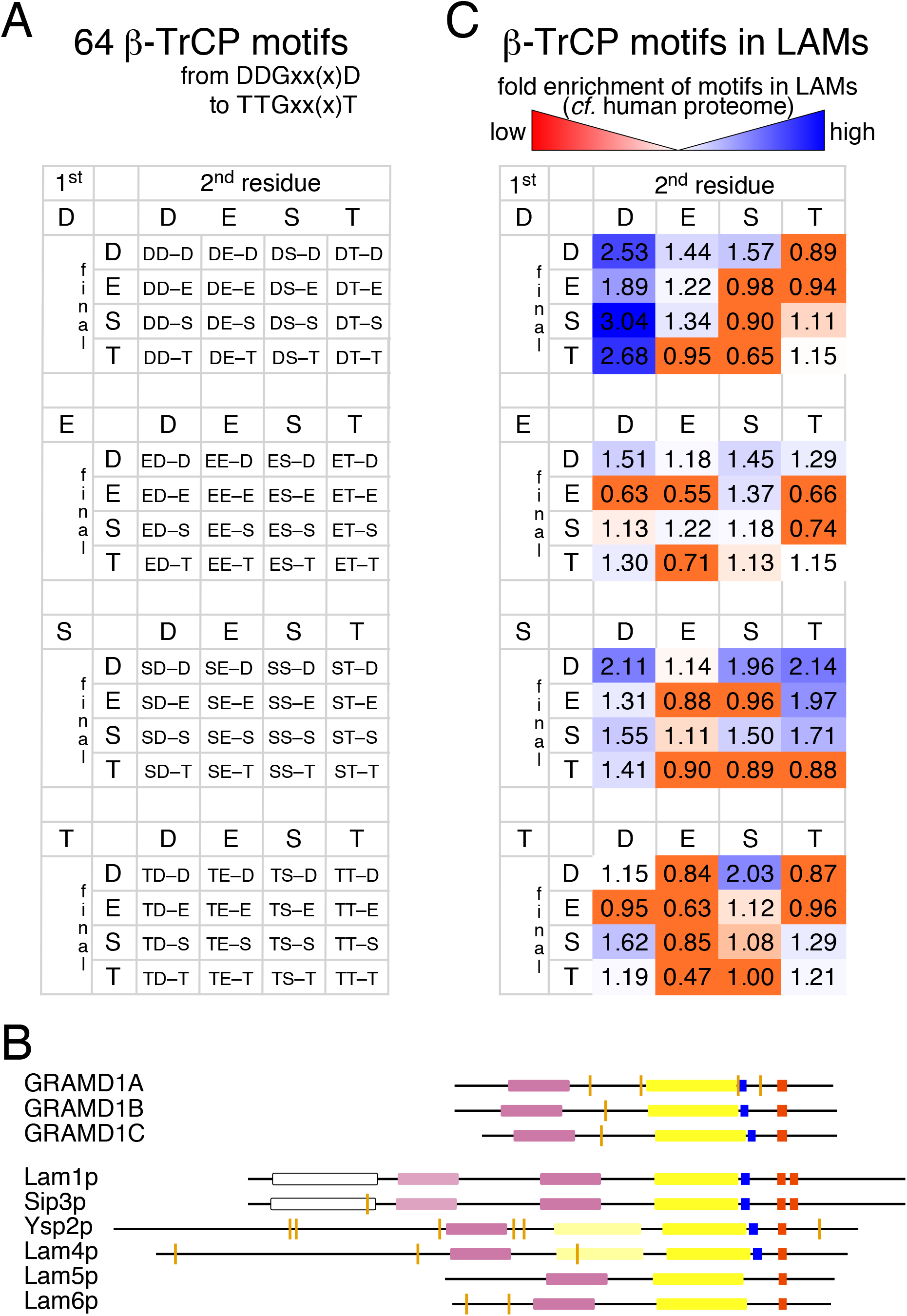
SCF-βTrCP Degrons in LAM proteins across evolution. **A.** 64 possible degron sequences of all possible combinations fitting the definition [DEST]-[DEST]-G-x(2,3)-[DEST], showing just the 1^st^, 2^nd^ and final residues. **B.** All instances of possible SCF-βTrCP degrons in human and yeast LAMs. Occurrences of [DEST]-[DEST]-G-x(2,3)-[DEST] shown by orange lines; other domains are the same as in Figure 1A/B, with additional PH domains in light pink and BAR domains in white. **C**. Fold enrichment of motifs in the LAM protein family (PF16016) over a random control protein set (the human proteome), layout as in A. Over-enriched motifs are coloured increasingly blue, and under-enriched motifs red. The most enriched motifs are DDGxx(x)-[DST].

**Supplementary Figure 3:**
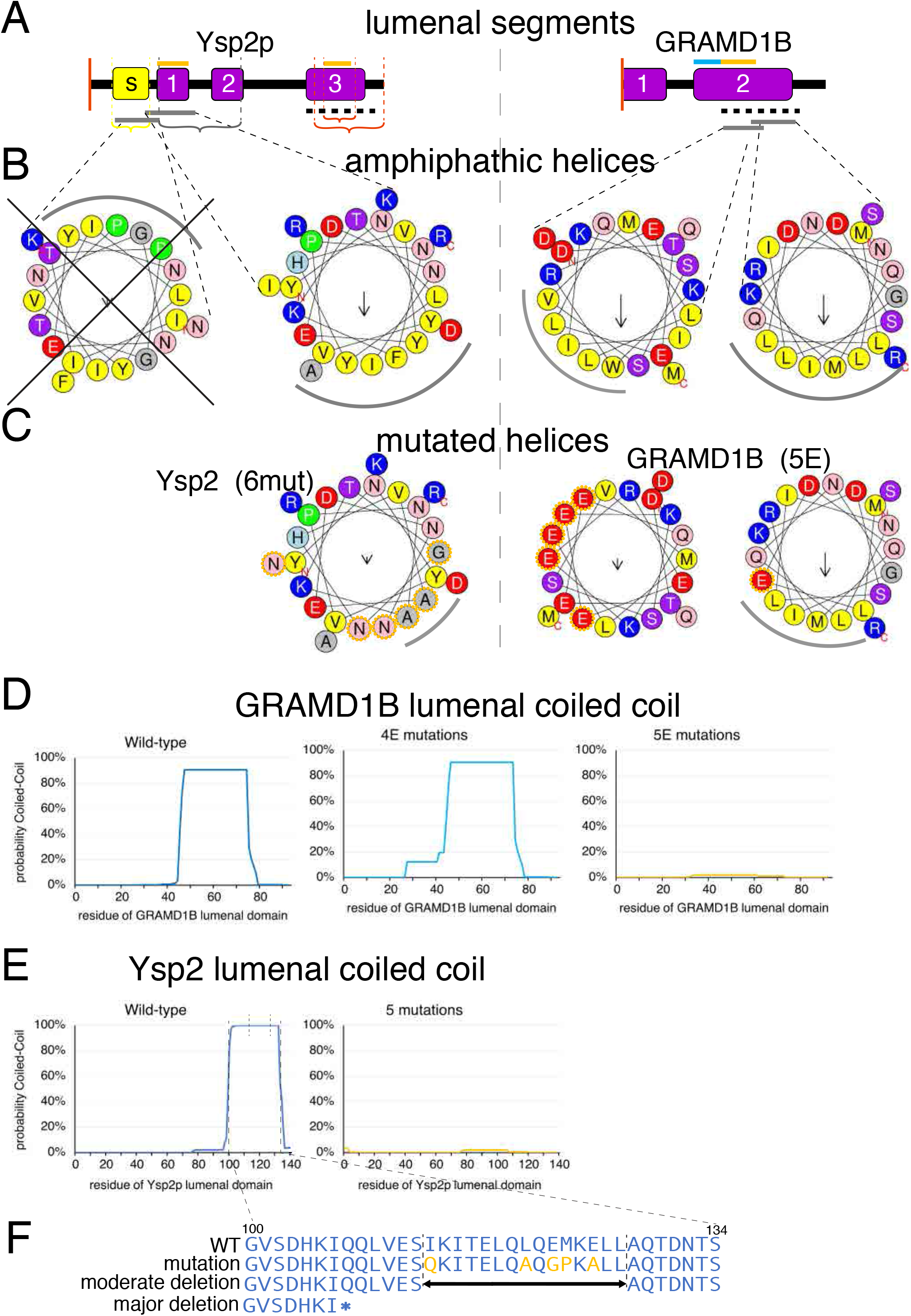
Details of helices in lumenal region of Ysp2 and GRAMD1B. **A.** Summary of elements in lumenal regions of Ysp2 and GRAMD1B. Overall form as in Figure 1D. Brackets indicate regions deleted in Ysp2p: sheet (yellow); amphipathic helix (grey); coiled-coil – complete and partial (both red). Lines above identify regions where series of single substitutions have been made: orange – substitutions designed to change function; light blue: control substitutions (GRAMD1B 4E only). Grey and dashed black lines below indicate amphipathic helices and coiled-coils (see below). Probability local distance difference test (pLDDT) averages for the predicted secondary structural elements are: Ysp2CT: sheet=65%, helix 1=57%, helix 2=63%, helix 3=75%; GRAMD1B-CT: helix 1 (extension of TMH)=79%, helix 2=86%. pLDDT=70% is the threshold between low and OK confidence. **B.** Hydrophobic faces (grey arcs) of stretches of ≥18 residues, when projected as a helical wheel, using Heliquest (Gautier et al., 2008). Regions that are all hydrophobic are excluded as being TMHs. Possible overlapping amphipathic helices are found in portions of Ysp2CT (left) and GRAMD1B (right). The first portion of Ysp2CT in unlikely to be an amphipathic helix (black cross superimposed) as the hydrophobic face contains two prolines, the hydrophobic moment is negative, and it is found in the region predicted to form β-strands. **C.** Substitutions in helices of Ysp2p and GRAMD1B that reduce amphipathicity. Substituted residues are highlighted by dotted orange lines. In Ysp2p, six substitutions of hydrophobic residues to asparagine (x3) alanine (x2) and glycine (x1) reduce the hydrophobic face to two alanines, preserving charge. In GRAMD1B, substitution of five hydrophobic residues to glutamate (5E) removes the first hydrophobic face entirely and reduces the second from 5 to 4 residues, and also changes overall charge. **D. and E.** Probability of coiled-coil formation by the C-terminus of GRAMD1B (D) and Ysp2p (E). Coils were predicted using the standard Lupas algorithm enacted in PCOILS at the Tuebingen Toolkit using the MTIDK weighted matrix and the 28 residue window (Gabler et al., 2020). For GRAMD1B (part D), the second and third graphs show respectively that control substitutions (4E) and test substitutions have the desired effect on coiled coil formation (Naito et al., 2019). For Ysp2p (part E), residues 100 to 134 are predicted to form a coiled coil that is lost with the substitutions. **F.** Constructs used to test the coiled-coil, showing the five substitutions (orange) and the 15 residues involved in the moderate deletion.

**Supplementary Figure 4:**
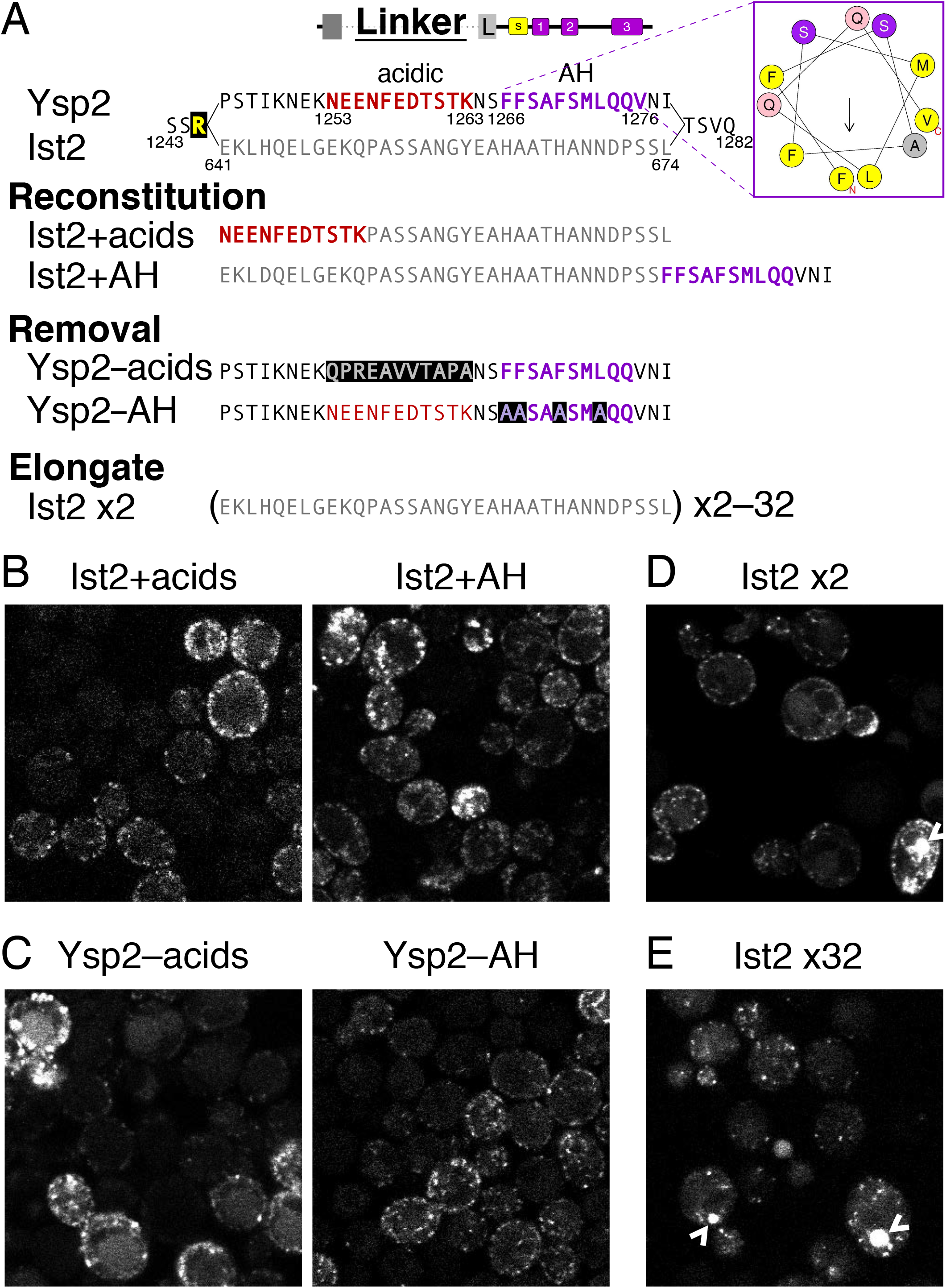
Details of experiments testing how the linker affects targeting by Ysp2CT. **A.** Sequences of different linkers used in constructs summarised in Figure 6G. Within the region of Ysp2 from 1243 to 1282 (with one extra arginine (yellow) added at the N-terminus after 1245 to allow cloning), 34 aa (1245-1278) were substituted with 34 aa from the linker in Ist2p (641-674). These and linkers with other changes as indicated were tested in constructs where the other regions were: neutralised PBR–generic TMH upstream, and the whole lumenal region downstream. Mutated residues are highlighted in black. **B.** Construct with Ysp2 linker as well as Ysp2 lumenal region. Arrows indicate internal fluorescent puncta, presumably located on the nuclear envelope. **C.** Two constructs that reconstitute Ysp2 elements into Ist2: either a stretch of acids from Ysp2p (1253-1263, red in part A), or a potential amphipathic helix from Ysp2 (1266-1276, see helical wheel, purple) was added into the Ist2 linker. Both additions restored punctate targeting. **D.** Two constructs removing elements from Ysp2. Either the stretch containing acids was replaced by electron-neutral residues from the linker of Lam5p (612-622: QPREAVVTAPA, light grey) or the potential amphipathic helix was mutated (four hydrophobics è alanine, light purple). Neither change prevented punctate targeting. **E and F.** Further constructs were based on elongation of the Ist2 linker, starting with a doubled linker (D: +34 aaè total 75 aa), and increasing two-fold to reach 32 copies of the linker (E:+1054 aaètotal 1095 aa). Large and bright intracellular dots (arrowheads) were seen rarely with 2 copies, becoming more common with increasing length of linker was the appearance of bright intracellular irregular shapes ≥0.5 µm in diameter (Supplementary Figure 4E), which appear to be artefactual aggregates. Similar objects were also commonly seen with wild-type Ysp2CT expressed from a much stronger promoter (*PHO5*).

**Supplementary Figure 5:**
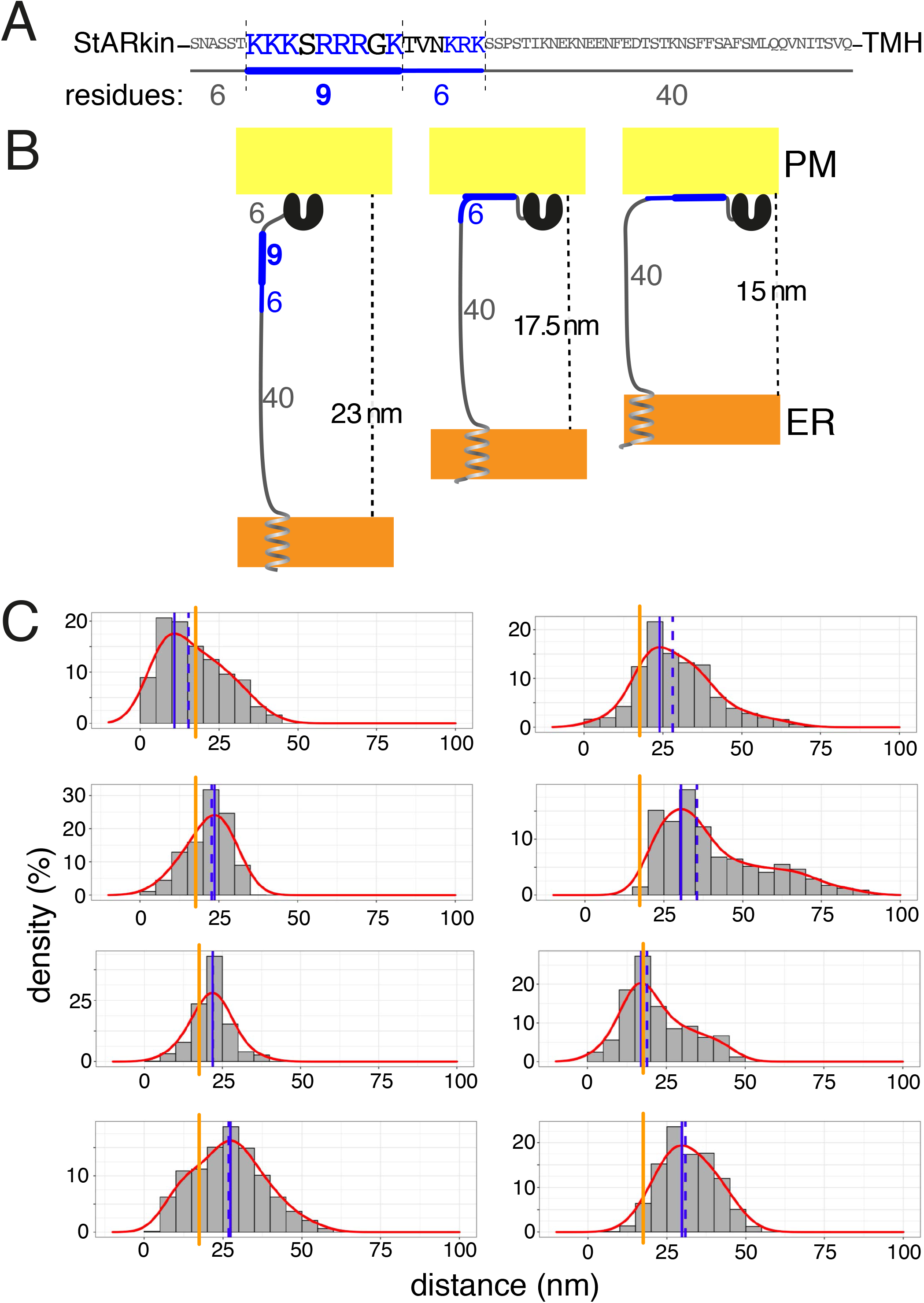
Histograms of ER-PM gaps at contact sites studied by Cryo Tomography. A. Sequence of the 61 residues in the Ysp2 linker between its second StARkin domain and its TMH. Background sequence is in grey, the PBR is shown in bigger font and blue for positively charged residues, with black for others in that region. The length of each segment of the PBR is shown below, along with a diagrammatic form in gray and blue lines. B. Diagram of three different conformations of the PBR envisaged in the text, allowing access of the StARkin domain to the PM either without any PBR contact (left), full engagement of the PBR (right) or partial engagement (middle), each with maximum width of the gap indicated. C. Each contact site imaged by electron tomography was analysed in 5×5 nm squares across the membrane plane (dimensions varying from 100 to 400 nm in each direction, membrane area from 16,000 to 76,000 nm^2^, corresponding to the ER-PM contact site imaged at a GFP-Ysp2CT spot. The graphs are annotated to indicate peak value (solid blue line) of the density function (red curve), medium (dashed blue line) and 17.5 nm (orange line). 17.5 nm corresponds to the maximum gap that can be crossed by 46 aa, thus allowing 9 PBR residues containing 7 lysine/arginines to interact with the PM, see Discussion.

